# Sleep-related thalamocortical spindles and delta oscillations are reduced during a ketamine-induced psychosis-relevant transition state

**DOI:** 10.1101/833459

**Authors:** A Mahdavi, Y Qin, A-S Aubry, D Cornec, S Kulikova, D Pinault

**Affiliations:** INSERM U1114, Neuropsychologie cognitive et physiopathologie de la schizophrénie, Strasbourg, France; Université de Strasbourg, Strasbourg, France; Fédération de Médecine Translationnelle de Strasbourg, FMTS, Faculté de médecine, Strasbourg, France; University of Freiburg, Bernstein Center Freiburg, Germany; Netherlands Institute for Neuroscience, Amsterdam, The Netherlands; National Research University Higher School of Economics, Perm, Russia

**Keywords:** Apomorphine, cell-to-network electrophysiology, clozapine, dizocilpine, midline thalamic nuclei, NMDA receptors, non-REM sleep, pentobarbital, physostigmine, quantitative EEG, somatosensory system, thalamic reticular nucleus

## Abstract

**Background:** In schizophrenia, sleep spindles are reduced, supporting the hypothesis that the thalamus and glutamate receptors play a crucial etio-pathophysiological role, whose underlying mechanisms remain unknown. We hypothesized that a reduced function of NMDA receptors is involved in the psychosis-related spindle deficit.

**Methods:** An electrophysiological multisite cell-to-network exploration was used to investigate, in sleeping rats, the effects of a ketamine-induced psychosis-relevant transition state in the sensorimotor and associative/cognitive thalamocortical (TC) systems.

**Results:** Under the control condition, spontaneously-occurring spindles (intra-frequency: 10-16 waves/s) and delta-frequency (1-4Hz) oscillations were recorded in the EEG of the frontoparietal cortex, in thalamic extracellular recordings (n=16), in dual juxtacellularly recorded GABAergic thalamic reticular nucleus (TRN) and glutamatergic TC neurons (n=8), and in intracellularly recorded TC neurons (n=8). The TRN cells rhythmically exhibited robust high-frequency bursts of action potentials (7 to 15 APs at 200-700 Hz). A single administration of low-dose ketamine fleetingly reduced TC spindles and delta oscillations, amplified ongoing gamma-(30-80Hz) and higher-frequency oscillations, and switched the firing pattern of both TC and TRN neurons from a burst mode to a single AP mode. Furthermore, ketamine strengthened the gamma-frequency band TRN-TC connectivity (n=11). The antipsychotic clozapine consistently prevented the ketamine effects on spindles, delta- and gamma-/higher-frequency TC oscillations (n=7).

**Conclusion:** The present findings support the hypothesis that NMDA receptor hypofunction is involved in the psychosis-related reduction in sleep spindles and delta oscillations. The ketamine-induced swift conversion (from burst to single APs) of ongoing TC-TRN activities may have involved both the ascending reticular activating system and the corticothalamic pathway.

**LAY ABSTRACT:** Schizophrenia is a chronic debilitating disease. Sleep disturbances associated with a reduction in spindles are observed as warning signs prior to the first psychotic episode. Every spindle is a short-lasting (~0.5 s) set of bioelectric sinusoidal waves at the frequency of 10-16 Hz generated within the thalamus. Sleep spindles, easily identifiable in a scalp electroencephalogram, occur hundreds of times during sleep and are implicated in cognition like memory processes. For this reason, spindles are seen as an electro-biomarker of the quality of sleep and cognitive performance. In patients at high risk of psychotic transition, the density (number/time unit) of spindles is reduced. The underlying mechanisms of this change are unknown. Glutamate-mediated neurotransmission in the thalamus plays a key role in the generation of spindles and the etiology of schizophrenia. Therefore, we tested the hypothesis that a reduced function of glutamate receptors at the thalamic level is involved in the psychosis-related reduction in spindles. Using cell-to-network neurophysiological methods in sleeping rats, we demonstrate that systemic administration of the NMDA glutamate receptor antagonist, ketamine, significantly decreases spindle density. This effect is consistently prevented by the widely used antipsychotic drug, clozapine. These original findings support the hypothesis of the involvement of a reduced function of NMDA glutamate receptors in the sleep spindle deficit observed in psychosis-related disorders. The present findings lay the foundation for the development of innovative therapies aimed at preventing psychotic, bipolar, and depressive disorders.

**HIGHLIGHTS:**

- Low-dose ketamine has a fast onset arousal promoting effect.
- Ketamine fleetingly reduces, in the first-/higher-order thalamocortical systems, sleep spindles and slow-waves, and amplifies gamma- and higher-frequency oscillations.
- Ketamine switches the firing pattern from a burst mode to a single action potential mode in both the glutamatergic thalamocortical neurons and the GABAergic thalamic reticular nucleus neurons.
- Ketamine strengthens the gamma-frequency band connectivity between thalamocortical and thalamic reticular nucleus neurons.
- The reference antipsychotic clozapine consistently prevents the ketamine effects.

## INTRODUCTION

Sleep abnormalities are detected not only during the early course of complex mental health diseases, such as schizophrenia (1–3) but also in individuals having a high-risk mental state for developing a transition to psychotic and bipolar disorders (4). Cortical EEG studies conducted in such patients have revealed a reduction in sleep spindles (5–9) and slow-wave activity (10). The underlying neural mechanisms are unknown. Sleep spindles have a thalamic origin with the GABAergic thalamic reticular nucleus (TRN) being a leading structure in their generation by exerting a powerful rhythmic inhibitory modulation of thalamocortical (TC) activities (11–13). The TRN is innervated by two major glutamatergic inputs, TC and layer VI corticothalamic (CT) axon collaterals, which mediate most of their excitatory effects through the activation of glutamate receptors (14–16). Importantly, layer VI CT axons innervate simultaneously TC and TRN neurons (17). The first-order TC systems are the specific, sensory and motor circuits, which receive cortical inputs only from layer VI CT neurons. The higher-order TC systems are the non-specific, associative/limbic/cognitive circuits, which receive cortical inputs from both layer V and layer VI CT neurons (18). In contrast to layer VI CT neurons, layer V CT neurons do not innervate the TRN. Of importance, the 3-neuron layer VI CT-TRN-TC circuit is robustly involved in the generation of sleep spindles (19, 20). There is accumulating evidence that dysfunction of thalamus-related systems is a core pathophysiological hallmark for psychosis-related disorders (21–25). NMDA receptors are also essential in the generation of thalamic spindles (26, 27), and a reduced function of these receptors is thought to play a critical role in the etio-pathophysiology of schizophrenia (28–31). Furthermore, the NMDA receptor antagonist ketamine models a transition to a psychosis-relevant state in both healthy humans (32–35) and rodents (36–39). Therefore, we hypothesized that a reduced function of NMDA receptors is implicated in the reduction of the density of sleep spindles recorded in patients having or about to have psychotic disorders. In an attempt to test this hypothesis, we investigated the effects of a low dose of ketamine on spindle oscillations recorded in first- and higher-order TC systems in the lightly anesthetized rat.

## METHODS AND MATERIALS

### Animals and drugs

Sixty-nine Wistar adult male rats (285-370 g) were used with procedures performed under the approval of the Ministère de l’Education Nationale, de l’Enseignement Supérieur et de la Recherche. Ketamine was provided from Merial (Lyon, France); clozapine, MK-801, apomorphine, and physostigmine, from Sigma-Aldrich (Saint-Quentin Fallavier, France), pentobarbital from Sanofi (Libourne, France), and Fentanyl from Janssen-CILAG (Issy-Les-Moulineaux, France).

### Surgery under general anesthesia

Deep general anesthesia was initiated with an intraperitoneal injection of pentobarbital (60 mg/kg). An additional dose (10-15 mg/kg) of pentobarbital was administered when necessary. Analgesia was achieved with a subcutaneous injection of fentanyl (10 μg/kg) every 30 minutes. The anesthesia depth was continuously monitored using an electrocardiogram, watching the rhythm and breathing, and measuring the withdrawal reflex. The rectal temperature was maintained at 36.5 °C (peroperative and protective hypothermia) using a thermoregulated pad. The trachea was cannulated and connected to a ventilator (50% air–50% O2, 60 breaths/min). The anesthesia lasted about 2 hours, the time necessary to perform the stereotaxic implantation of the electrodes (40).

### Cortical EEG recordings under analgesic pentobarbital-induced sedation

In 28 rats, a recording silver wire (diameter: 200 μm) sheathed with Teflon was implanted in the parietal bone over the primary somatosensory cortex (from bregma: 2.3 mm posterior and 5 mm lateral). At the end of the surgery, the rectal temperature was set to and maintained at 37.5°C. The analgesic pentobarbital-induced sedation was initiated about 2 h after the induction of the deep anesthesia and maintained by a continuous intravenous infusion of the following regimen (average quantity given per kg and per hour): Pentobarbital (4.2 ± 0.1 mg), fentanyl (2.4 ± 0.2 μg), and glucose (48.7 ± 1.2 mg). To help maintain a stable mechanical ventilation and to block muscle tone and tremors, a neuromuscular blocking agent was used (d-tubocurarine chloride: 0.64 ± 0.04 mg/kg/h). The cortical EEG and heart rate were under continuous monitoring to adjust the infusion rate to maintain the sedation.

### Thalamic cell-to-network recordings under analgesic pentobarbital-induced sedation

The recording-labeling glass micropipettes were filled with a saline solution (potassium acetate, 0.5 M) and a neuronal tracer (Neurobiotin, 1.5%). Three series of experiments were performed: 1) The first (16 rats) was designed to perform, along with the cortical EEG, extracellular (field potential and single/multiunit) recordings in first- and higher-order thalamic nuclei. The regions of interest were stereotaxically (41) located behind the bregma (2.3 to 3.6 mm posterior). 2) In the second series (8 rats), paired juxtacellular TC and TRN recordings were performed in the somatosensory system. The diameter of the micropipette tip was about 1 μm (15-30 MOhms) (42). 3) The third (11 rats) was designed to record intracellularly TC neurons. The diameter of the micropipette tip was inferior to 1 μm (30–70 MOhms). The extracellular and juxtacellular signals (0.1-6000 Hz), and the intracellular signal (0-6000 Hz) were acquired using a low-noise differential amplifier (DPA-2FL, npi electronic, GmbH) and an intracellular recording amplifier (NeuroData IR-283; Cygnus Technology Inc.), respectively. All signals were sampled at 20 kHz 16-bit (Digidata 1440A with pCLAMP10 Software, Molecular Devices). At the end of the recording session, the target neurons were individually labeled with Neurobiotin using the extra- or juxtacellular nano-iontophoresis technique (42) to identify formally both the recording site and the structure of the recorded neurons (Fig1B). Then the animal was humanely killed with an intravenous overdose of pentobarbital, transcardially perfused with a fixative containing 4% paraformaldehyde in 10 mM phosphate buffer saline, and the brain tissue was processed using standard histological techniques for anatomical documentation.

**Figure 1:**
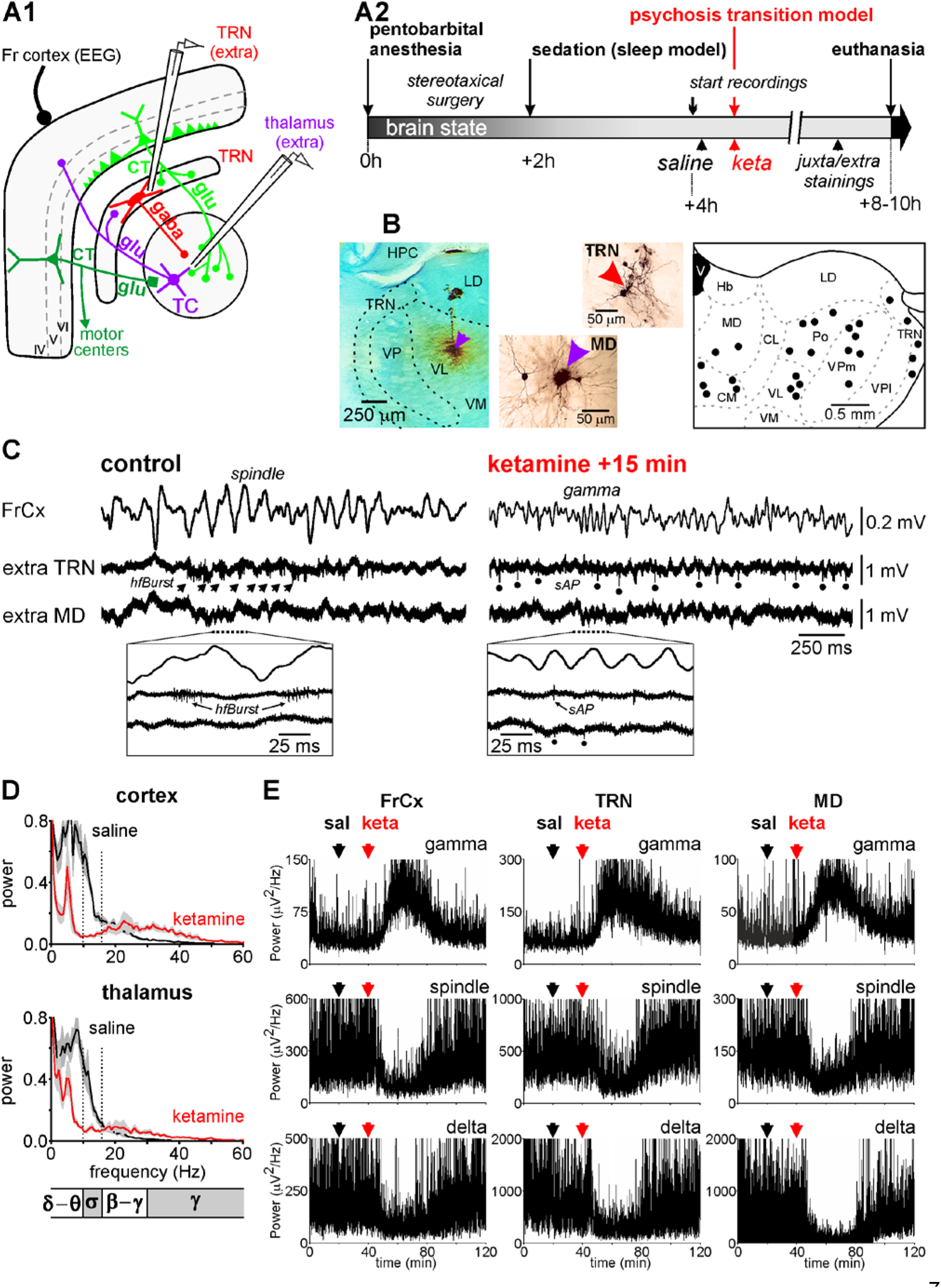
Ketamine reduces sleep oscillations in the thalamocortical systems. **(A1)** Experimental design showing the location of the two glass micropipettes designed to record the extracellular activities in the thalamic reticular nucleus (TRN) and in a dorsal thalamic nucleus along with the EEG of the frontal cortex. The hodology of the 4-neuron CT-TRN-TC circuit is also shown. The corticothalamic (CT) and thalamocortical (TC) neurons are glutamatergic while the TRN neuron is GABAergic. The cortical inputs of the first-order thalamic nuclei (like the ventral posterior, VP (somatosensory), and the ventral lateral, VL (motor)) originate from layer VI whereas the cortical inputs of the higher-order thalamic nuclei (like the posterior group, Po, or intralaminar/midline nuclei) originate from layers V and VI. In contrast to the layer V CT neurons, the layer VI CT neurons do innervate the TRN. The intrathalamic innervation pattern of layer VI CT neurons is regional whereas that of the layer V CT neurons, these latter CT neurons targeting only high-order thalamic nuclei, is more punctual. The layer V CT axon, which does not innervate the TRN, is a branch of the corticofugal main axon that targets the lower motor centers (brainstem and spinal cord). **(A2)** Design timeline illustrating the principal steps of the experiment. The color code of the brain state is dark gray for anesthesia, light gray for sedation and dark for death. **(B)** The left microphotograph shows, at low-magnification, the track left by the electrode and the extracellular labeling of the neurons located at and close to the recording site (here in the VL); the middle microphotograph shows, at higher-magnification, the recording site in the thalamic medial dorsal nucleus (MD, indicated by the arrowhead) and the somatodendritic complex of a couple of MD neurons; the left microphotograph shows the recording site with a few neural elements labeled in the TRN. On the right is presented, into a coronal plane, a mapping of the recording sites (black dots) into the TRN and the dorsal thalamic nuclei. The coronal plane represents a block of brain of about 3.6 mm thick posterior to the bregma (from −2.3 to −3.6 mm) in which recordings were performed. **(C)** Under the saline (control) condition, the cortex, the TRN and the dorsal thalamic nuclei exhibit a synchronized state, characterized by the occurrence of low-frequency (1-16 Hz) oscillations, including spindles. The extracellular TRN recordings can contain high-frequency (200-700 Hz) bursts of APs (hfBurst, indicated by arrows). The framed expanded trace shows a couple of hfBursts associated with TC spindle waves. Under the ketamine condition, the TC system displays a more desynchronized state, characterized by the prominent occurrence of fast activities (>16 Hz), which include gamma-frequency oscillations. And the TRN cell fires more in the single AP (sAP) mode than in the hfBurst mode. Extracellular sAPs are indicated by the dots. Below, the expanded trace reveals sAPs associated with TC gamma waves. Single APs are also identifiable (indicated by dots) in the extracellular recording of the MD under the ketamine condition. **(D)** Spectral analysis of the cortical EEG (top) and of the thalamic extracellular activities (bottom) recorded under the saline then the ketamine conditions. Each value is a grand average (±SEM) from 6 rats, each rat being its control (per value: 23 epochs of 2 s/rat (hamming, resolution: 0.5 Hz)). In each chart, the part delimited by 2 dotted lines indicates the sigma-frequency band, which corresponds predominantly to spindles. **(E)** Time course of the power of, from top to bottom, gamma oscillations, spindles, and delta oscillations recorded simultaneously in the frontal cortex (FrCx), the TRN and in the medial dorsal (MD) nucleus before and after subcutaneous administrations of saline and ketamine (at 20 and 40 min, respectively).

### Data analysis

Analysis software packages Clampfit v10 (Molecular Devices) and SciWorks v10 (Datawave Technologies) were used. Spectral analysis of EEG and network oscillations was performed with fast Fourier transformation (FFT, 2-Hz resolution). The power of baseline activity was analyzed in 4 frequency bands: delta-(1–4 Hz), sigma-(10–16 Hz, spindles), gamma-(30–80 Hz), and higher-(81–200 Hz) frequency oscillations. For each band, the total power was the sum of all FFT values. In single-unit juxtacellular recordings, single action potentials (APs) were detected using a voltage threshold and an inter-AP interval superior to 10 ms. High-frequency bursts (hfBursts) were identified based on a voltage threshold and an inter-AP interval inferior to 4 ms. A TC or TRN burst had a minimum of 1 inter-AP interval. Inter-AP time and autocorrelogram histograms, and the density (number per minute) of single APs and hfBursts were computed. To apprehend the time relationship between the network or cellular gamma waves and the cellular firing of a single TC or TRN neuron, a 25-55 Hz filter was used to make gamma waves detectable, to create a peri-event time histogram of the TC or TRN firings. Standard inter-AP interval (resolution 1 ms) histograms were computed. Each drug effect was measured relative to the vehicle condition with each rat being its control. Statistical significance of the observed effects was evaluated with the Student’s paired t-test (significant when P≤0.05).

## RESULTS

### Ketamine reduces thalamocortical spindles and delta-frequency oscillations

The arousal promoting effect of a single low-dose (2.5 mg/kg) of ketamine has been well documented in free-behaving rats (37, 38, 43). Moreover, under the ketamine condition, not a single sleep episode was observed during the time dedicated to sleep (S1). To understand how ketamine could influence ongoing sleep oscillations, cell-to-network recordings were performed in the TC system of pentobarbital sedated rats, a rodent model of slow-wave sleep with spindles (44). Our recordings started about 2 hours after the onset of the infusion of the pentobarbital containing regimen (Fig1A2), that is when the on-line spectral analysis revealed a stationary amount of spindles and slower oscillations (FigS2, Fig1C), which were qualitatively similar to those recorded during the natural non-REM sleep (FigS1B2). Multisite extracellular recordings were performed in the TRN, in midline, posterior, and ventral thalamic nuclei. The extracellular recordings could contain low-amplitude, single-/multi-unit firings (Fig1C). Notably, TRN cells exhibited rhythmic hfBursts in relation to the sleep TC oscillations (Fig1C). From ~5 minutes after a subcutaneous administration of ketamine (2.5 mg/kg), the pattern of the cortical and thalamic baseline sleep activities was dramatically reduced in amplitude, supplanted by a more desynchronized pattern (Fig1C). Indeed, ketamine significantly decreased the power (synchronization index) of the spindles and delta oscillations (Fig1D,E, FigS3). It also decreased the amount of theta-frequency (5-9 Hz) oscillations (Fig1D), a peak of CT theta activity that is a hallmark of drowsiness (45). Concomitantly, ketamine significantly increased the power of ongoing gamma- and higher-frequency oscillations. The ketamine effects, observed in all recorded regions (n≥4 rats/region; Fig2), were transient (peaking at 15-20 minutes) with partial recovery at 60-80 minutes after the administration (Fig1E). In contrast to drugs modulating dopaminergic and cholinergic transmitter systems, dizocilpine (MK-801), a more specific NMDA receptor antagonist, well mimicked the ketamine effects on spindles and higher-frequency oscillations (S4).

**Figure 2:**
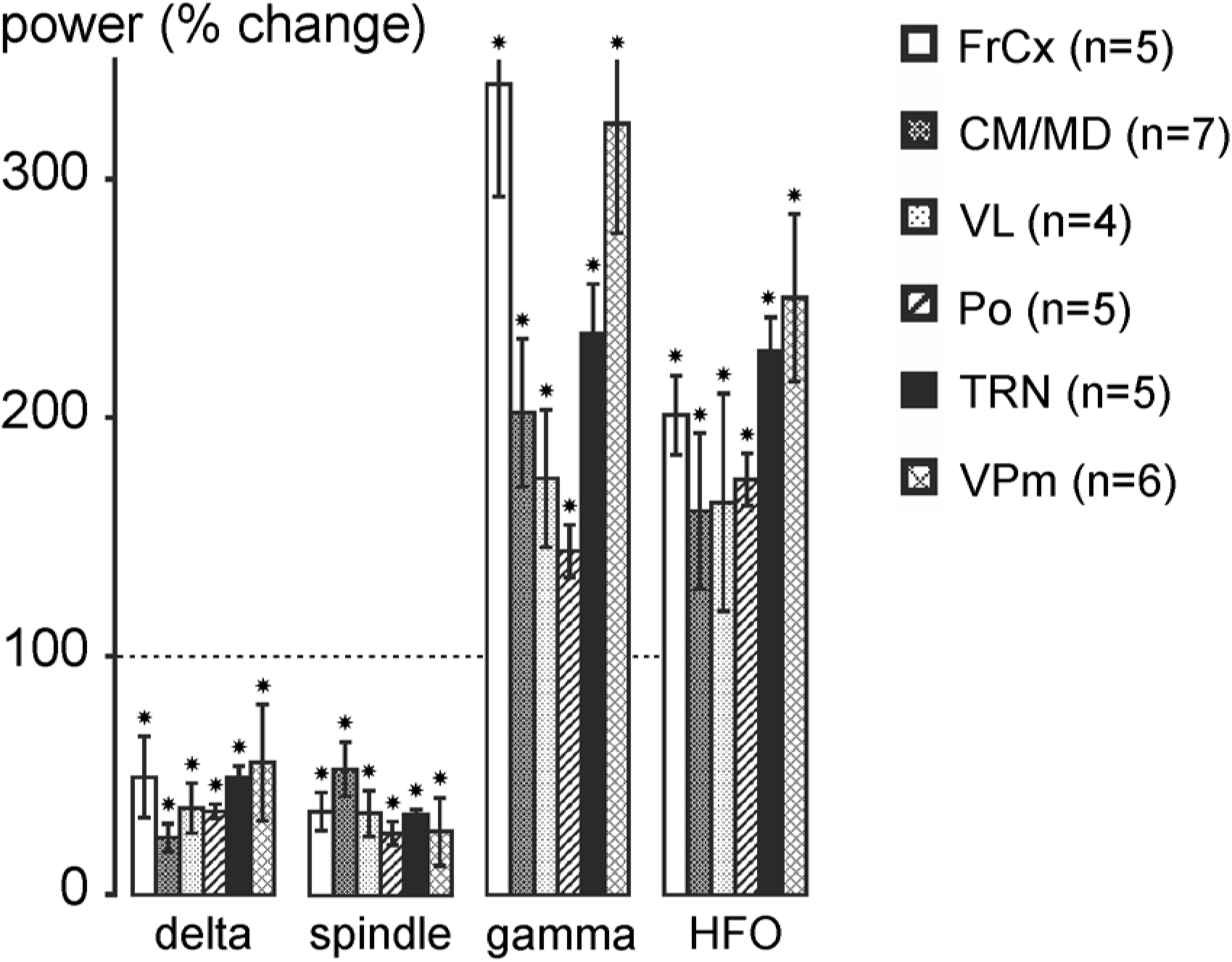
Ketamine reduces delta-frequency oscillations and spindles and increases gamma- and higher-frequency oscillations. The histogram shows the ketamine-induced percent changes (±SEM, relative to the saline condition, each rat being its control; post-ketamine: 20 to 30 minutes) in power of delta oscillations, spindles, gamma- and higher-frequency oscillations recorded in the frontal cortex, in the TRN and in first-order (VPm, VL) and in higher-order (CM/MD, Po) thalamic nuclei. Number of rats given in parentheses. Paired t-test relative to saline condition (star when p < 0.05). For abbreviations, see figure 1 legend.

### Ketamine switches the firing pattern of thalamic relay and reticular neurons from the burst mode to the tonic mode

In the following, all data are from the somatosensory system as it contains less than 1% of local-circuit neurons (46) and its 3-neuron layer VI CT-TRN-TC circuit, common to first- and higher-order thalamic nuclei, is the leading circuit in the generation of spindles. The location of the recording sites was identified based on electrophysiological and anatomical features (FigS5). From 11 extracellular thalamic recordings, 6 (from 6 rats) contained at least two TC units that were detectable using an automated spike sorting procedure. Five out of 8 dual juxtacellular TC-TRN recordings (5 rats) had a duration long enough for data analyses under control and ketamine conditions, and 8 out of 15 TC cells met the intracellular requirements (44).

#### Thalamic relay neurons

Low-amplitude APs of one or more neurons were often visible in extracellular recordings (Fig1C, FigS6A). During the sedation, the TC units presented an irregular firing pattern consisting in hfBursts and single APs. It was extremely rare to see series of rhythmic hfBursts at the spindle frequency, suggesting that most of the TC spindle oscillations were subthreshold (44), as demonstrated by the dual juxtacellular TRN-TC recordings (Fig3A1) and by the intracellular recordings of TC neurons (Fig3D). From ~5 minutes after the ketamine administration (16 TC units from 6 rats), the density of hfBursts significantly decreased whereas that of single APs increased for at least 60 minutes (FigS6B). The spike sorting method may, however, not be precise and reliable as the amplitude and shape of the APs might not be stationary over time (47). For instance, in TC hfBursts, the AP amplitude became progressively smaller (FigS6A). Therefore, to better validate the ketamine effects observed in the extracellular TC recordings, we performed dual juxtacellular recordings of thalamic relay and reticular neurons. The juxtacellular single-unit recording-labeling technique allows the formal identification of the recorded neuron (Fig3A1,B1,C1) (42). Ketamine, transiently and significantly, decreased the density of TC hfBursts and increased that of single APs (Fig3A1,A2 and B3). However, the decrease in the hfBurst density was ~50%, meaning that AP bursts still occurred under the ketamine condition. Embedded in the irregular tonic AP trains, a lot of them were doublets and triplets, whose intra-frequency was lower (inter-AP interval peak at 5-6 ms, Fig4A1,A2) than that of typical hfBursts (interval peak at 2-3 ms, Fig4A1). A partial recovery was noticeable 60-80 minutes after the ketamine administration (FigS6B, and Fig3B3). Of importance, ketamine increased the firing frequency band of TC neurons from, on average, 5-20 Hz (10.8±2.9 Hz, N=5 from 5 rats) to 15-30 Hz (21.7±3.5 Hz, N=5) (Fig3E). Furthermore, in one of the experiments, designed to record in the posterior group (equivalent to the pulvinar in humans) of the thalamus, 2 nearby (100 μm apart) TC cells were simultaneously recorded in the juxtacellular configuration (FigS7). Ketamine consistently augmented their firing frequency band in a similar way (from 0-10 Hz to 0-35 Hz).

**Figure 3:**
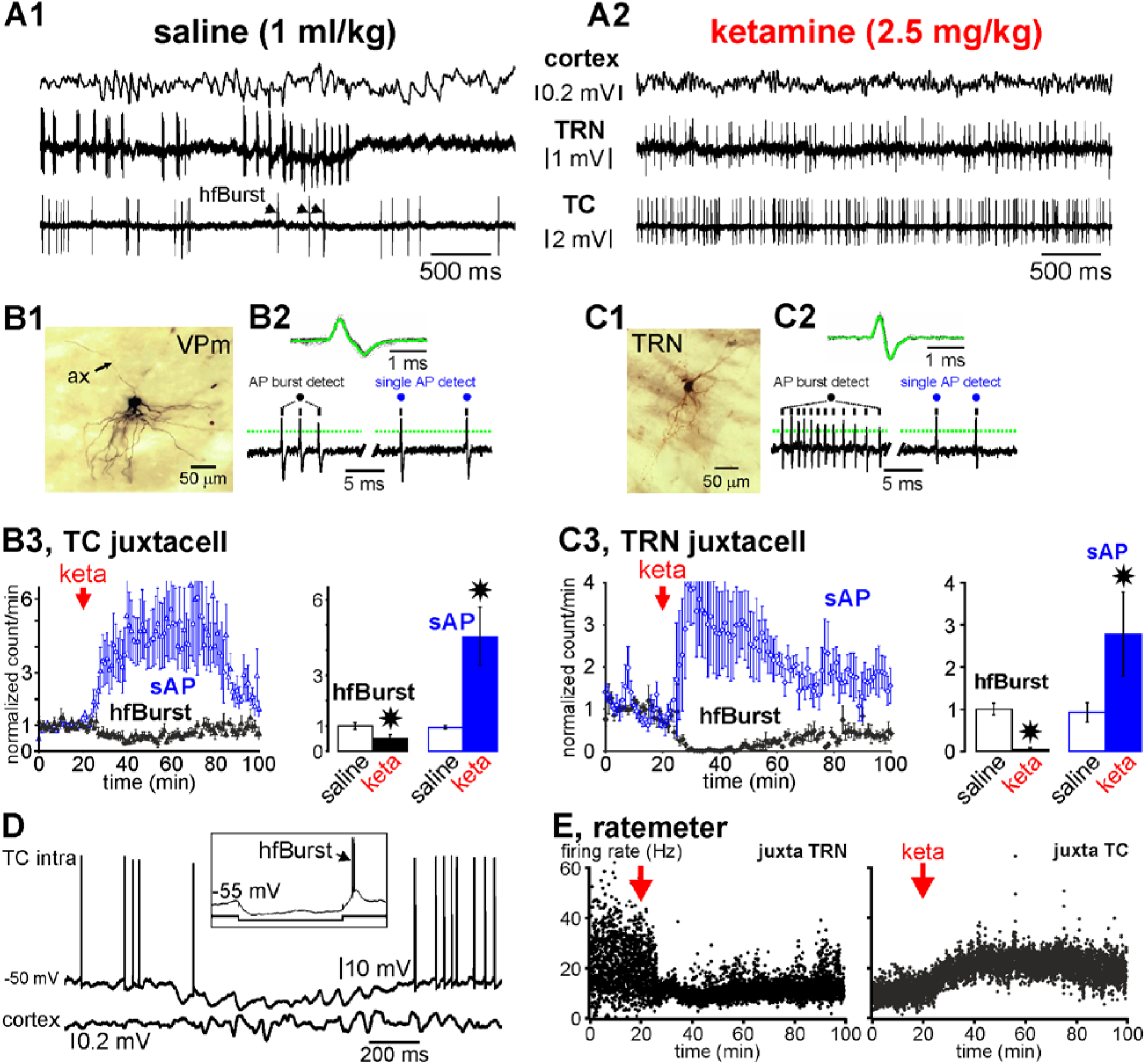
Ketamine switches the firing pattern from a burst mode to a single action potential mode in thalamic relay (glutamatergic) and reticular (GABAergic) neurons. **(A1, A2)** Typical simultaneous recordings of the cortex (EEG), and of two single TRN and TC neurons (juxtacellular configuration) of the somatosensory system. Under the saline (A1, control) condition, the cortex displays a synchronized state, characterized by the occurrence of medium-voltage (>0.1 mV) low-frequency (1-16 Hz) oscillations, the TRN cell exhibits a typical series of rhythmic robust high-frequency bursts of action potentials (hfBursts, 300-500 APs/s), and the TC neuron exhibits single action potentials (sAPs) and, during the TRN burst series, a few bursts. A few minutes after the systemic administration of ketamine (A2, here: +20 minutes), the cortex displays a more desynchronized state, characterized by the prominent occurrence of lower voltage (<0.1 mV) and faster activities (>16 Hz), which include gamma-frequency oscillations. Under the ketamine condition, both the TC and the TRN cells exhibit much more sAPs than hfBursts. **(B1-B3)** Data from juxtacellularly recorded TC neurons. **(B1)** Photomicrography of parts of the somatodendritic complex and of the main axon (ax) of a juxtacellularly recorded and labeled (with Neurobiotin) TC neuron of the somatosensory thalamus. **(B2, top)** Average and superimposition of 50 action potentials. **(B2, below)**: Detection (from a voltage threshold, indicated by a dotted line) of a typical hfBurst of 3 APs and of 2 successive single APs. **(B3)** The density (number per minute, ±SEM, 5 TC cells from 5 rats) of hfBursts and of sAPs under the saline and ketamine conditions. Paired t-test (star when p<0.05). **(C1-C3)** Data from juxtacellularly recorded TRN neurons. **(C1)** Photomicrography of part of the somatodendritic complex of a juxtacellularly recorded and labeled (with Neurobiotin) TRN cell. **(C2, top)** Average and superimposition of 50 APs. **(C2, below)**: Detection (from a voltage threshold, indicated by a dotted line) of a typical hfBurst of 12 APs and of 2 successive single APs. **(C3)** The density (number per minute, ±SEM, 5 TRN cells from 5 rats) of hfBursts and of sAPs under the saline and ketamine conditions. Paired t-test (star when p<0.05). **(D)** Representative trace of an intracellularly recorded TC neuron showing the occurrence of subthreshold oscillations, including spindle-frequency rhythmic waves, which are concomitant with a synchronized EEG state in the related cortex. Note that the subthreshold oscillations occur during the through of a long-lasting hyperpolarization. In the frame is shown the occurrence of a low-threshold potential topped by a high-frequency burst of APs (hfBurst) at the offset of a 200-ms hyperpolarizing pulse. **(E)** Ratemeter of simultaneously juxtacellularly recorded TRN and TC neurons under saline then ketamine conditions. Each dot is the average (n = 5 neurons from 5 rats) of the number of inter-AP intervals per second.

**Figure 4:**
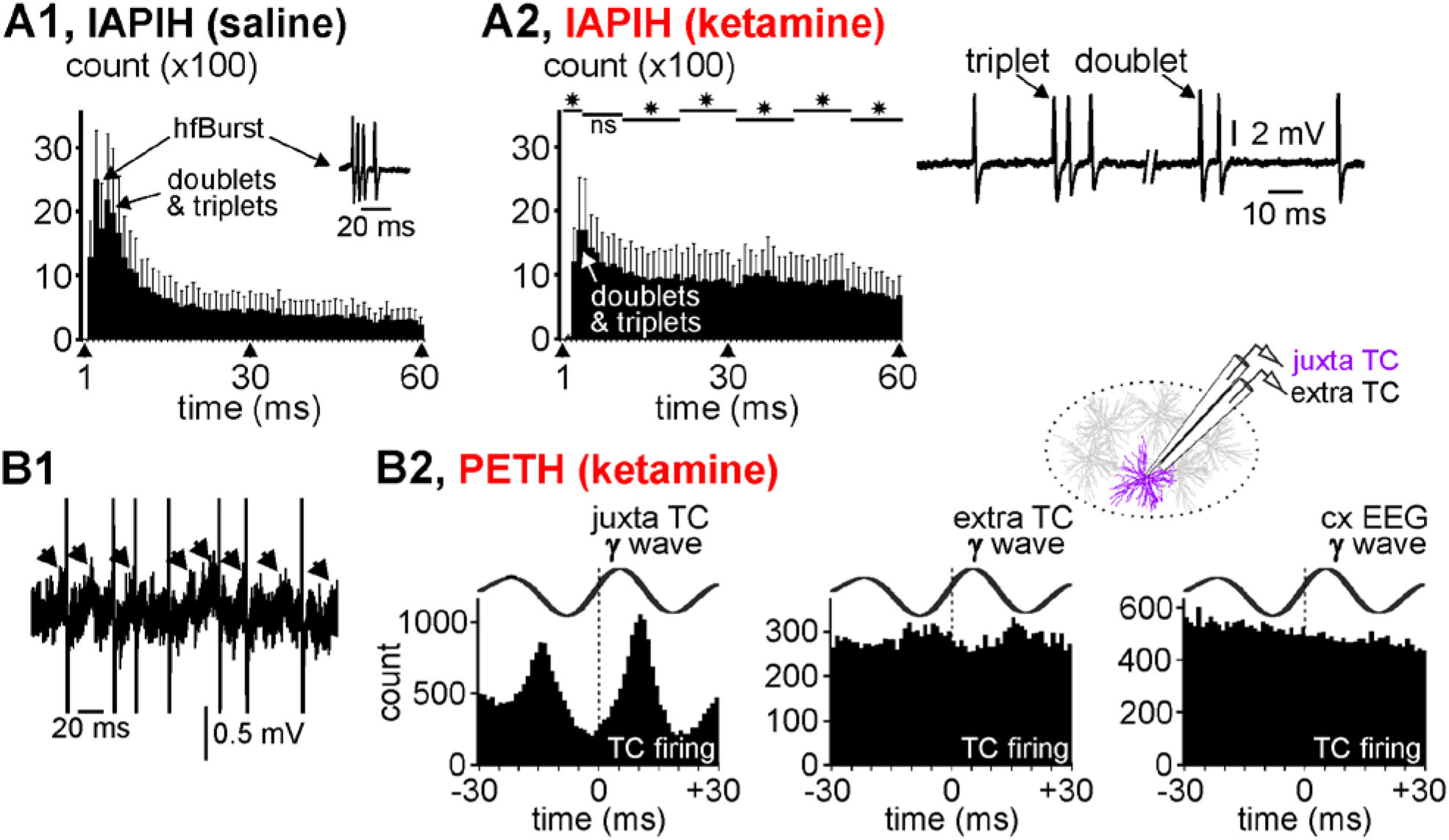
Thalamocortical firing related to gamma-frequency oscillations. **(A1, A2)** Averaged cumulated inter-AP interval histograms (IAPIH) from 5 juxtacellularly recorded TC cells (from 5 rats) under the control (A1) then the ketamine (A2) conditions (keta +15-25 min). Note the ketamine-induced diminution in the number of the short-lasting IAPIs, especially those composing high-frequency bursts of APs (IAPI = 2-10 ms). A typical TC hfBurst (IAPI at 2-3 ms) is shown in the control histogram. The remaining bursts were slower (IAPI at 5-6 ms) and shorter (e.g, especially doublets and triplets, like those shown on the right). Star when significant (paired t-test, p <0.05). **(B1)** A typical short-lasting trace of a juxtacellularly recorded TC cell showing low-amplitude gamma-frequency oscillations in the field potential and the AP occurrence at some cycles of the gamma oscillation. Each arrow indicates a juxtacellular gamma wave. The APs are truncated. **(B2)** Peri-event (gamma wave) time histogram (1-ms resolution) of the TC firing (cumulative count) under the ketamine condition (5 TC cells from 5 rats). Every gamma wave (juxta TC, extra TC (inter-tip distance = 100 μm, see drawing), and cxEEG) is an average of 100 filtered (25-55 Hz) individual gamma (γ) waves. Time “0” corresponds to the time at which gamma waves were detected.

Curiously, under the ketamine condition, the mean firing frequency of TC neurons (<30 Hz) was lower than the network gamma-frequency oscillations (frequency at maximal power: 33.6±1.1 Hz, n=7), raising the question whether or not TC single APs were related to the juxta- and extracellular gamma oscillations. In an attempt to address this question, firstly we looked at the raw juxtacellular recordings, in which we notice that TC neurons did not emit an AP at every wave of the gamma oscillations (Fig4B1), suggesting that the juxtacellular field potential variations reflected more membrane potential oscillations than APs. Secondly, a substantial number of single APs was phase-related to both the juxtacellular and the extracellular (100 μm apart) gamma waves (FigS8, Fig4B2). However, the temporal link was stronger with the juxtacellular (cellular activity) than the extracellular (nearby network activity) wave. In contrast to layer-organized cortical structures, the weak relation between the juxtacellular APs and the extracellular gamma waves seen in the somatosensory thalamus might have been due to an anarchic overlap of the current sinks and sources generated by the neural activities. On the other hand, there was no apparent relation between the TC firing and the cortical gamma waves (Fig4B2), which is not surprising as the EEG integrates the activities of interweaved large-scale networks.

#### Thalamic reticular neurons

During sedation, all TRN cells recorded with extra- or juxtacellular recording methods exhibited sequences of rhythmic hfBursts (Fig1C, Fig3A1). The burst sequence naturally recurred at a low frequency (<1 Hz) (FigS2B1 and FigS5B), during which rhythmic hfBursts occurred at the sigma (spindle)- and lower-frequency bands, including the delta band. The rhythmic character of spindle burst patterns was identifiable with an autocorrelation histogram (Fig5A1). In TRN neurons, such sustained rhythmic burst activity involves the activation of NMDA receptors (27). From ~5 minutes after a single ketamine administration, all juxtacellularly recorded TRN cells suddenly and transiently switched their ongoing rhythmic burst firing pattern to a sustained tonic, single AP firing pattern (Fig3A1,A2,C3). Furthermore, ketamine decreased their firing frequency band from 0-60 Hz (16.5±2.5 Hz) to 5-25 Hz (8.1±1.8 Hz; n=5 from 5 rats) (Fig3E). Remarkably and significantly, the single AP density increased whereas the hfBurst density decreased (Fig3C3, Fig5A2, B1-B2). In the inter-AP interval histogram, the first peak at 2-4 ms, a marker of hfBursts, disappeared almost completely. Under the ketamine condition, the first peak (3-6 ms) reflects longer inter-AP intervals which, like in TC neurons, are the signature of doublets and triplets embedded in the irregular tonic AP trains (Fig5B2).

**Figure 5:**
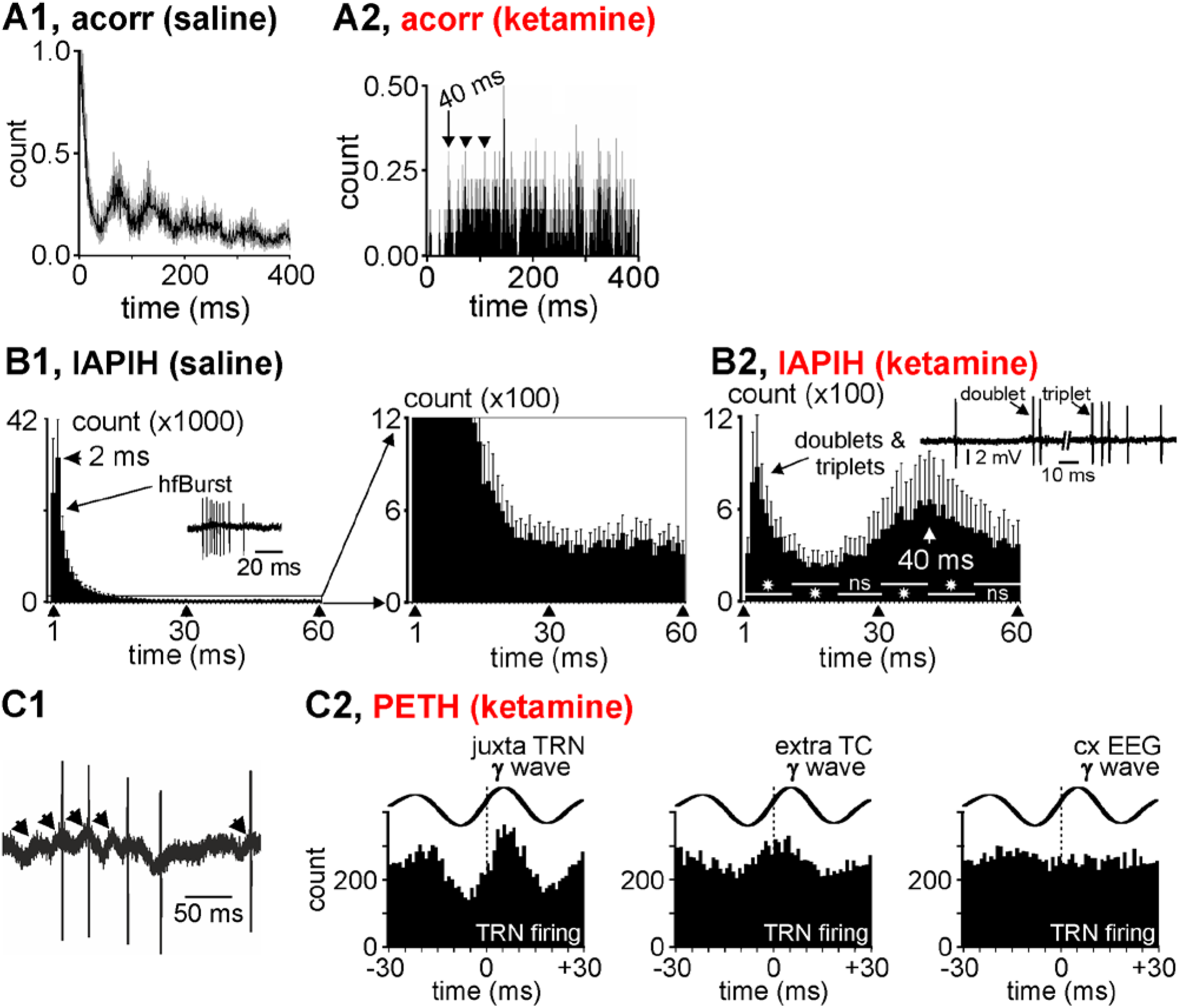
Thalamic reticular nucleus firing related to gamma-frequency oscillations. **(A1, A2)** Averaged autocorrelograms (acorr, resolution: 1 ms, from ten 2-s traces ±SEM in grey) of the firing of a representative juxtacellularly-recorded TRN neuron under the saline (A1) and ketamine (A2) conditions. Under the ketamine condition, 2 to 3 successive APs (arrow and arrowheads have a probability of about 0.20-0.25 (relative to the reference AP at time 0 ms) to be separated by an interval of 40 ms. The units of the Y axis are normalized counts. **(B1, B2)** Averaged cumulated inter-AP interval histograms (IAPIH) from 5 juxtacellularly recorded TRN cells (from 5 rats) under the control (B1) then the ketamine (B2) conditions (keta +15-25 min). In (B1), are shown the control (saline) IAPIH in full (left) and partial (right) Y scales. Note the ketamine-induced dramatic diminution in the number of the short-lasting IAPIs, especially those composing high-frequency bursts of APs (IAPI = 2-10 ms). A typical TRN hfBurst is shown in the control histogram. The remaining bursts were slower (increase in IAPIs) and shorter (e.g, especially doublets and triplets, like those shown). Star when significant (paired t-test, p <0.05). **(C1)** A typical short-lasting trace of a juxtacellularly recorded TRN cell showing low-amplitude gamma-frequency oscillations in the field potential and that the AP occurrence at some cycles of the gamma oscillation. Each arrow indicates a juxtacellular gamma wave. **(C2)** Peri-event (gamma wave) time histogram of the TRN firing (cumulative count) under the ketamine condition (5 TRN cells from 5 rats). Every gamma wave (juxta TRN, extra TC, and cxEEG) is an average of 100 filtered (25-55 Hz) individual gamma (γ) waves. Time “0” corresponds to the time at which gamma waves were detected.

Interestingly, the interval histogram reveals a second peak at ~40 ms. Furthermore, the autocorrelation histogram shows that the first 2 to 3 successive APs had a probability of about 0.20-0.25 (relative to the reference AP at time 0) to be separated by an interval of 30-50 ms (Fig5A2). We predicted that the 30-50-ms peak represents a marker of juxtacellular gamma oscillations. Indeed, when looking closely at the juxtacellular recordings, it is obvious that the TRN cells fired at a certain proportion of gamma waves during their positive-going component (Fig5C1), meaning that the juxtacellular oscillations reflected threshold/suprathreshold and subthreshold membrane potential gamma oscillations. This observation is supported by a peri-gamma wave time histogram of the AP distribution (Fig5C2), which shows that the probability of firing reached a maximum at (virtually 0 ms) the positive-going component of the gamma wave. Furthermore, a substantial number of TRN APs was also phase-related to gamma waves recorded extracellularly in the related somatosensory thalamic nucleus, suggesting a certain degree of functional connectivity. Moreover, using the partial correlation coefficient (S9), the strength of the gamma-frequency band TRN-TC connectivity was significantly increased by ketamine (Fig6). On the other hand, in the same way as TC neurons, there was no apparent relation between the TRN firing and the cortical gamma waves (Fig5C2).

**Figure 6:**
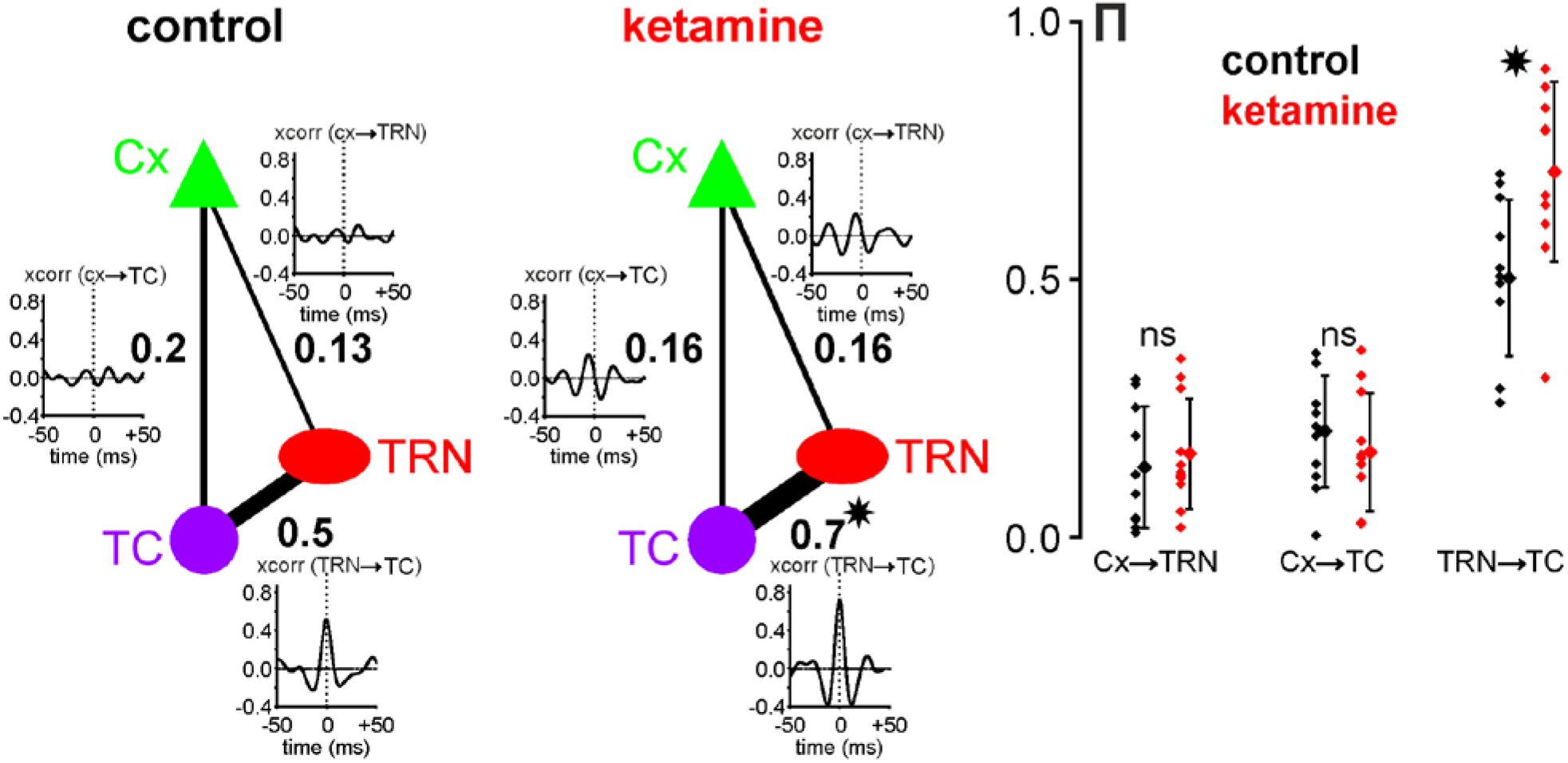
Ketamine strengthens the functional gamma TRN-TC connectivity. Direct interaction strength between two different sites is given by a partial correlation coefficient written (bold font) next to the edge connecting these sites. The plots presented next to these edges are cross-correlograms of 400ms-epochs from signals recorded at the corresponding sites and filtered in the gamma-range (25-55Hz). Under control condition, the strength of gamma interactions between TRN and TC sites was more than twice higher than for CT-TRN and CT-TC interactions, which was also reflected by a high peak in the TRN-TC cross-correlogram. When ketamine was applied, the strength of TRN-TC gamma interactions was significantly increased (paired t-test, p<0.001), resulting in a higher partial correlation coefficient and a higher peak in the average cross-correlogram. Although after ketamine application, correlations between CT and TRN and between CT and TC were higher in the cross-correlograms, the strength of CT-TRN and CT-TC interactions given by partial correlation coefficients did not change significantly (paired t-test, p>0.4). The plot on the right shows distributions of partial correlation coefficients *⊓* for CT→TRN, CT→TC and TRN→TC gamma interactions in all experiments (N=11) under both control (black) and ketamine (red) conditions. (*) indicates significant difference revealed with a paired t-test with p<0.001; ns, non-significant.

#### Clozapine prevents the ketamine effects

Clozapine is one of the most effective antipsychotic drugs against treatment-resistant schizophrenia (48). Its clinical effects are thought to be related to interactions with a variety of receptors, including the glutamatergic receptors and more specifically NMDA receptors via the glycine site (49–51). Also, clozapine is well-known to modulate sleep spindles (52), possibly due to the activation of GABAergic TRN neurons via a specific action on D4 dopamine receptors (53), which would exert a tonic influence on the TRN activity (54). Therefore, it was interesting to probe whether a single systemic administration of clozapine could prevent the ketamine effects on TC oscillations. To address this issue, clozapine was subcutaneously administered at a dose (5 mg/kg) that durably decreases the power of spontaneously-occurring cortical gamma oscillations in the naturally-behaving rat (55) 20 or 120 minutes before the ketamine challenges. In all rats (n=7), clozapine consistently prevented the ketamine peak (at ~15-20 min) effect on spindles, delta- and gamma-/higher-frequency oscillations (FigS10 and Fig7). When administered alone, clozapine significantly increased the power of delta oscillations (Fig7).

**Fig 7:**
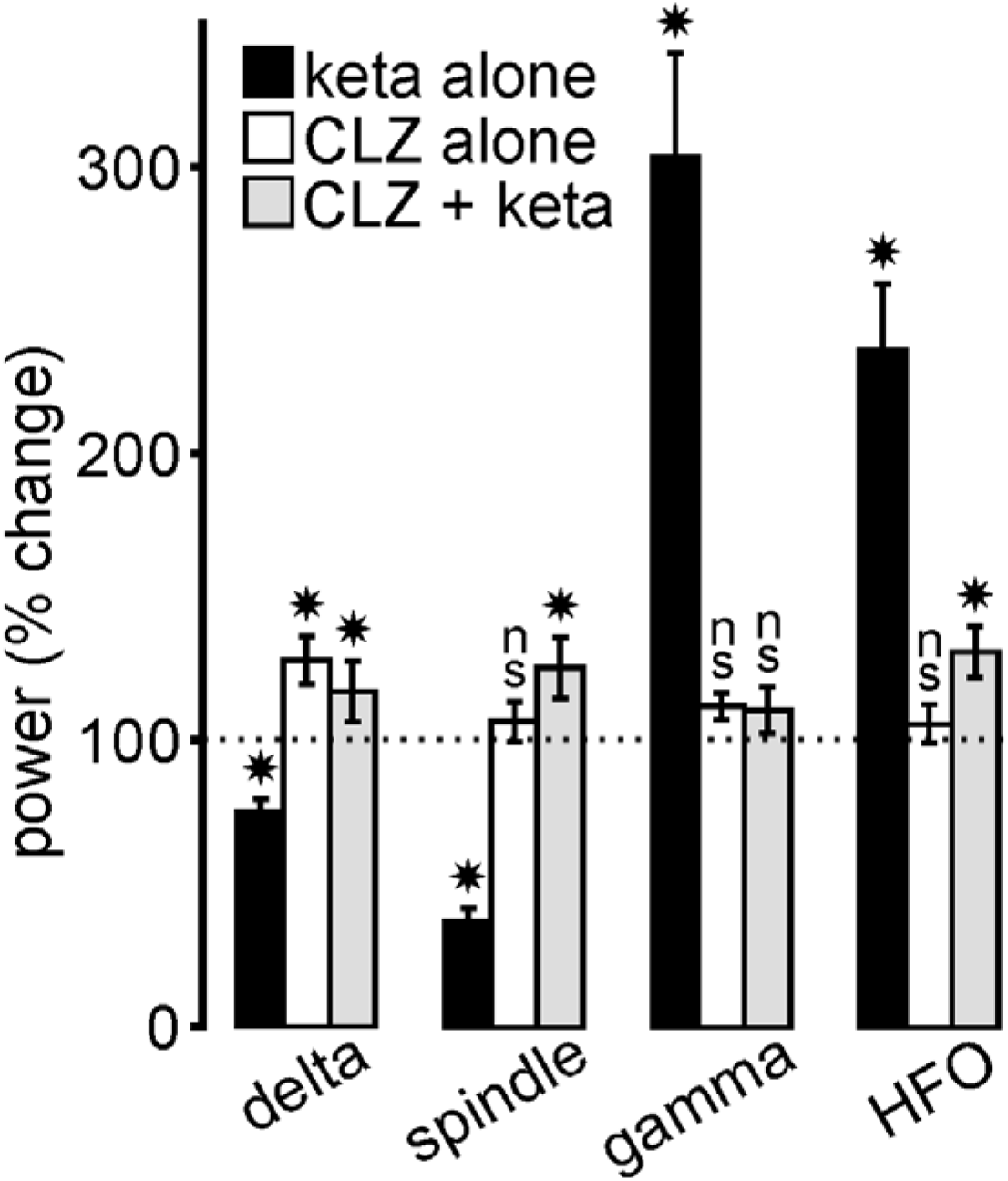
Clozapine (CLZ) prevents the ketamine effects. The histogram illustrates the drug-induced percent changes (mean±SEM; relative to their respective vehicle (saline for ketamine, saline/HCl 0.1N for clozapine) condition (100%, indicated by dotted line); 5 rats/condition) in power of all frequency bands in the cortical EEG. Student paired t-test: (*) p< 0.05; ns, not significant. In the CLZ + keta condition, ketamine (2.5 mg/kg) was administered 20 minutes after the CLZ (5 mg/kg) administration.

## DISCUSSION

These novel results support the hypothesis of the involvement of a reduced function of NMDA receptors in the deficit in sleep spindles and delta oscillations observed in psychosis-related disorders. The translation of these findings to human diseased condition provides a strong foundation for the use of NMDA-related medications against various forms of psychoses.

### The arousal promoting effect of low-dose ketamine

In free-behaving rats, ketamine dose-dependently increased the level of wakefulness associated with a psychosis-relevant behavior (37, 38), which delayed the sleep onset latency (43). Clinical investigation showed that patients with psychosis have difficulties initiating sleep (56). Abnormal levels of arousal may be a predictor of psychotic disorders (57, 58). Here, it is further shown that, in the sedated rat, ketamine elicited a fleeting arousal reaction, at least in the TC-TRN system, which is electrophysiologically reminiscent of REM sleep, a brain state considered as a natural model of psychosis (59–63). Moreover, the NMDA receptor hypofunction-related increase in gamma-/higher-frequency oscillations observed in sedated rats is also recorded during the natural REM sleep (64). So, we interpret the ketamine-induced desynchronized state as uncharacteristic REM sleep or a pathological persistent UP state (Fig8). During the ketamine-induced pathological UP state, expected to occur within diverse cortical and subcortical structures (38), cortical and thalamic neurons would be more depolarized than during the DOWN state to generate more threshold (for AP initiation) and supra-threshold membrane potential oscillations (65). In thalamic neurons, the burst mode is a reliable hallmark of sleep oscillations, every hfBurst occurring at the top of a low-threshold Ca^++^ potential mediated by the activation of T-type channels, which are de-inactivated via membrane hyperpolarization (<-60 mV) (66). Both the synaptic interactions between TC and TRN neurons and the intrinsic pacemaker properties of TRN cells are well-known to play leading roles in the generation of thalamic spindles (11, 13). Under the ketamine condition, the substantial increase in the single AP density suggests that the membrane potential of TC and TRN neurons was more often depolarized. This is supported first by the occurrence of gamma oscillations and single AP firing in our juxtacellular TC and TRN recordings and, second, by an increase in the gamma band TRN-TC connectivity. The single AP mode is usually recorded when T-type Ca^++^ channels are inactivated via membrane depolarization (>-60 mV) (67). Disruption of the CaV3.3 Ca^++^ channel, which encodes the low-threshold T channels (68), may be involved in the etio-pathophysiology of schizophrenia (69).

**Figure 8.**
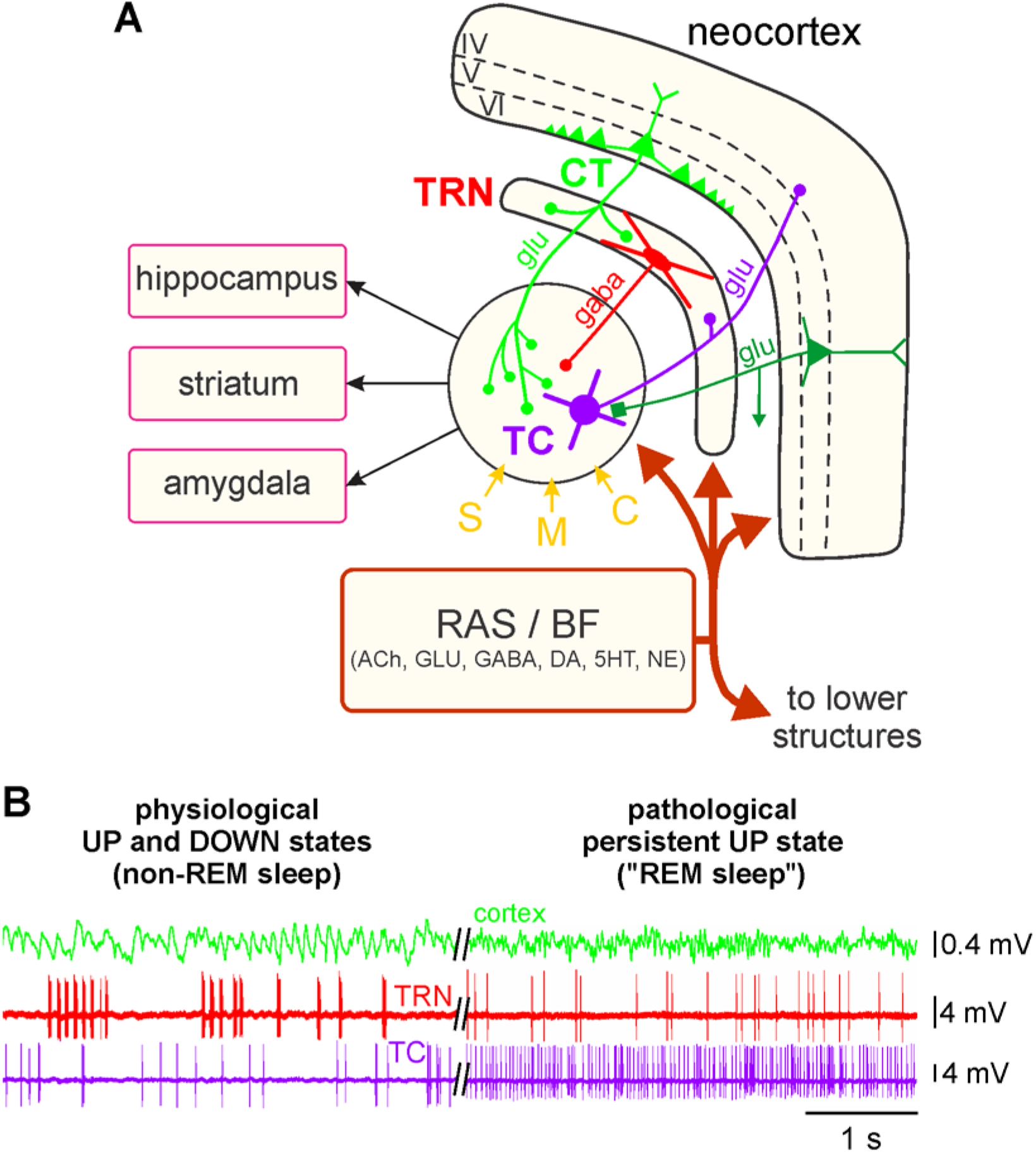
Theoretical prediction of the ketamine action in both the ascending reticular activating system and the corticothalamic pathway. **(A)** Simplified drawing of the hodology of the 4-neuron CT-TRN-TC circuit, which is considered as being the upper part of the ascending reticular activating system (RAS). See main text and figure 1 legend for detailed description of the circuit, which receives sensory (S), motor (M) and cognitive/associative (C) inputs. It is important to specify that the layer VI CT neurons outnumber by a factor of about 10 the TC neurons. The first- and higher-order thalamic nuclei are under the neuromodulatory influence of the various inputs from the ascending RSA and the basal forebrain (BF). **(B, left)** Physiological UP and DOWN states: During the non-REM sleep, the TC system displays principally a synchronized state, characterized by the occurrence of delta oscillations and spindles; the TRN cell exhibits mainly rhythmic (at the delta-, theta- and spindle-frequency bands) hfBursts of action potentials. The synchronized state includes two sub-states, UP and DOWN, which are usually associated with active and quiescent cellular firings, respectively. **(B, right)** Pathological persistent UP state: This ketamine-induced persistent UP state is assumed to be an abnormal REM sleep. After a single systemic administration of a subanesthetizing low-dose of ketamine, the TC system displays a more desynchronized state (peak effect at about +15-20 minutes) characterized by the prominent occurrence of lower voltage and faster activities (>16 Hz), which include beta-, gamma- and higher-frequency oscillations. Under the ketamine condition, both the TC and the TRN neurons exhibit a persistent irregular and tonic firing containing more single APs than hfBursts. ACh, acetylcholine; GLU, glutamate; 5HT, serotonin; DA, dopamine; NE, norepinephrine.

### Contribution of the corticothalamic pathway

In the thalamus, ketamine would act principally on both the glutamatergic TC and the GABAergic TRN neurons. How did ketamine convert the firing from burst to the tonic mode in both TC and TRN neurons? During sleep, sustained hyperpolarization would be the result of either excess inhibition or disfacilitation. Under the ketamine condition, a likely effect would be a sustained excitation of these two types of neurons by common afferent input. In addition to the influence of neuromodulatory inputs (see below), the CT pathway seems an excellent candidate (70, 71). Such a scenario would involve the disinhibition of CT neurons, leading them to sustained network gamma hyperactivity. Such CT hyperactivity can enhance thalamic gamma oscillations (14, 72). This puts forward the hypothesis of greater sensitivity of cortical GABAergic neurons to NMDA receptor antagonists (73, 74), which would be responsible for the disinhibition of CT neurons. If this scenario is true, the ketamine-induced hyperactivation of layer VI CT neurons could promote the gamma-frequency pacemaker properties of the GABAergic TRN cells (75, 76).

NMDA receptors are more critical for the CT-mediated excitation of TRN than TC neurons (26). Furthermore, the long-lasting kinetics of NMDA receptors in the GABAergic TRN neurons are essential to promote rhythmic Ca^++^-mediated burst firing, which then cyclically hyperpolarizes the postsynaptic TC neurons through the activation of GABA receptors. Importantly, in TRN neurons, the NMDA-mediated effects of CT transmission can work across a wide range of voltages so as the voltage-dependent blockade by Mg^++^ is incomplete and that NMDA receptors can be activated by synaptically released glutamate even in the absence of AMPA receptor-mediated activation (26). Thus, because CT neurons outnumber by a factor of ~10 TC neurons (77), the ketamine NMDA-mediated effects are expected to be stronger on TRN than on TC neurons in reducing burst activity, which, in fact, was indeed observed in the present study (Fig3B3,C3). The ketamine-induced increase in thalamic gamma power may have been due to a specific blockade of NMDA receptors involving specifically the NR2B subunit (64), which is present in the TRN (78). The NMDA receptor hypofunction-related spindle reduction and gamma increase in the TRN-TC system may help to understand the increased TC connectivity correlated with spindle deficits in schizophrenia (33).

### Contribution of the ascending reticular activating system and basal forebrain

The likely mechanisms underlying the effects of ketamine remain debatable as it acts in all brain structures and at many receptors (79–81). The ketamine-induced acute arousal effect may involve, among many others, cholinergic, monoaminergic, and orexinergic arousal systems (43, 82, 83). In the present study, the fact that a single low-dose of ketamine simultaneously affected, in an opposite manner, spindle-/delta-frequency and gamma-/higher-frequency TC oscillations is reminiscent of the seminal finding of Moruzzi and Magoun (84). Indeed, these pioneering investigators demonstrated that electrical stimulation of the reticular formation, a complex set of interconnected circuits within the brainstem, evokes in the TC system a switch of the EEG pattern from a synchronized to a desynchronized state, an effect interpreted as an EEG arousal reaction. Thus, the present findings give further support to the hypothesis of a dysregulation of the ascending reticular activating system, which includes the pedunculopontine nucleus, the basal forebrain, and the thalamus, in the etio-pathophysiology of psychotic disorders (83, 85–87). Also, in the observed ketamine effects, we should not exclude a contribution of the ascending GABAergic pathways (originating from the brainstem, midbrain, ventral tegmental area, zona incerta, basal ganglia, and from the basal forebrain), which play a critical role in promoting TC activation, arousal and REM sleep (88, 89). In the present study, interestingly, physostigmine, known to promote REM sleep (90) and a cortical EEG arousal (91, 92), exerted a ketamine-like effect on the firing pattern of TRN neurons (FigS4c). The atypical antipsychotic clozapine consistently prevented the foremost ketamine-induced acute effects on sleep oscillations. The fact that, in contrast to ketamine, clozapine alone exerted no change in the cortical gamma power suggests that ketamine and clozapine exerted their action via distinct neural/molecular targets, which does not discredit the hypothesis of a dysregulation of the reticular activating system (83, 85–87), the TC system being nothing but its downstream part (Fig8).

### Conclusions and significance

The present preclinical investigation with its limitations (S11) demonstrates that the acute effects of ketamine result in fast onset arousal promoting effect, suggesting that it acts like a rapid-acting inducer of REM sleep-associated cognitive processes, which is reminiscent of its ability to induce hallucinatory and delusional symptoms (93–97). Low-dose ketamine not only disturbs brain rhythms, but also disrupts attention-related sensorimotor and cognitive processes (34, 98, 99), supporting the notion that schizophrenia is a cognitive disorder with psychosis as a subsequent consequence (100–102). The ketamine-induced changes in rodent EEG oscillations are reminiscent of those observed in at-risk mental state individuals (103, 104) and during the first episode of schizophrenia (105, 106). Taken together, the present findings support more strongly the whole brain-networks hypothesis than the isolated brain circuit theory of schizophrenia (107).

The neural mechanisms underlying the ketamine-induced fleeting arousal effect may be, in part, those responsible for the initial stage of the rapid-acting antidepressant action of ketamine in patients with drug-resistant major depressive disorders (108–110), leading us to think that the ketamine effects are state-dependent. In addition, the present results suggest that the combined sleep and ketamine models have some predictive validity for the first-stage development of innovative therapies against psychotic, bipolar, and depressive disorders.

## ACKNOWLEDGEMENTS

The present work was supported by INSERM, the French National Institute of Health and Medical Research (Institut National de la Santé et de la Recherche Médicale, 2013-), l’Université de Strasbourg, Unistra (2013-), and Neurex. This project has been funded with support from the NeuroTime Erasmus+ program of the European Commission (2015-2020: AM and YQ). This publication reflects the views only of the authors, and the Commission cannot be held responsible for any use which may be made of the information contained therein. ASA is a graduate student from the Euridol Graduate School of Pain. Data of the present study were presented in 2018 at both the FENS Forum (Berlin) and the SFN meeting (San Diego). The authors thank Yoland Smith and Martin Deschênes for critical reading of the manuscript.

## AUTHORS’ CONTRIBUTION

AM, YQ, SK, DP: Design, data acquisition & analysis, and writing; ASA: Data acquisition & analysis; DC: Animal well-being, surgery and technical aspects.

