## Supplemental Material for "Sleep-related thalamocortical spindles and delta oscillations are reduced during a ketamine-induced psychosis-relevant transition state"

### S1: Ketamine prevents the occurrence of natural sleep episodes

To determine whether a single sub-anesthetizing (assumed psychotomimetic) low-dose (2.5 mg/kg) of ketamine was able, in an acute fashion, to disturb the occurrence of natural sleep episodes, cortical EEG recordings were performed, in free-behaving rats, in the middle of the light phase (09:30 am-04:30 pm), that is, when the rat naturally falls asleep.

**Methods and Materials:** Six Wistar adult male rats (3-6-month-old, 285-370 g) were used. They were housed and kept under controlled environmental conditions (temperature: 22±1dC; humidity: 55±10%; 12/12 h light/dark cycle; lights on at 7:00 am) with food and water available *ad libitum*. Under surgical anesthesia (see main text), two recording silver wires (diameter: 200 μm) sheathed with teflon were stereotaxically (1) implanted in the right and left parietal bones over the bilateral frontoparietal cortex (from bregma: 2.3 mm posterior and 4.5 mm lateral). Only the section (0.02 $mm^2$) of each wire was in contact with the inner plate of the bone. Two screws were fixed in the frontal bone for ground connection and in the bone covering the cerebellum for the reference. The electrodes were connected to a connector fixed to the skull with dental cement. Recording sessions began after a 1-week recovery. After one week of training to the experimental conditions, the bilateral parietal cortical EEG (bandpass: 0.1-800 Hz; sampling rate: 10 kHz) was acquired using ultralow noise amplifiers (AI 402, x50; Axon Instruments) for 80 minutes after a subcutaneous administration of saline or ketamine (2.5 mg/kg). During the recording sessions, the behavior was rated every minute as immobile or passive (e.g, ball-like position, eyes closed or open), active (exploration, eating, chewing, motion, grooming), whisking, etc.

**Results:** After a 1-week habituation to the 80-min recording conditions, on average three (3.0±0.2, 18 recording sessions) sleep episodes were recorded under the pre-control (no injection) or the control (following a subcutaneous administration of saline, 1 ml/kg) condition. During sleep, the rats were usually in a ball-like position with their eyes closed. The onset of the sleep episodes usually started with non-rapid eye movement (non-REM) sleep accompanied in the EEG with intermixed K complexes, prominent delta- (1-4 Hz), theta- (5-9 Hz) and spindle- or sigma- frequency (10-16 Hz) oscillations (FigS1A1,B2). After a single subcutaneous administration of ketamine (2.5 mg/kg), not a single slow-wave sleep episode was observed during the recording session in all ketamine-treated rats (n = 5, 15 recording sessions; FigS1A2 and histogram). They were abnormally hyperactive with stereotypies and ataxia, and their cortex transiently displayed a remarkable increase in the power of beta- (17-29 Hz), gamma- (30-80 Hz) and higher-frequency (81-200 Hz) oscillations (FigS1A2,B3) as previously described (2, 3). One week later, the rats had a normal wake-to-sleep behavior as the number of slow-wave sleep episodes (n = 3.3±0.3, 11 recording sessions under the saline condition) was well comparable to that of the first control (saline) condition (FigS1, histogram). These findings show that ketamine prevented the occurrence of natural sleep during at least the 80-min recording sessions.

1. Paxinos G, Watson C (1998): *The rat brain in stereotaxic coordinates*. Fourth Edition ed.: Academic Press.
2. Pinault D (2008): N-methyl d-aspartate receptor antagonists ketamine and MK-801 induce wake-related aberrant gamma oscillations in the rat neocortex. *Biol Psychiatry*. 63:730-735.
3. Hakami T, Jones NC, Tolmacheva EA, Gaudias J, Chaumont J, Salzberg M, et al. (2009): NMDA receptor hypofunction leads to generalized and persistent aberrant gamma oscillations independent of hyperlocomotion and the state of consciousness. *PLoS One*. 4:e6755.

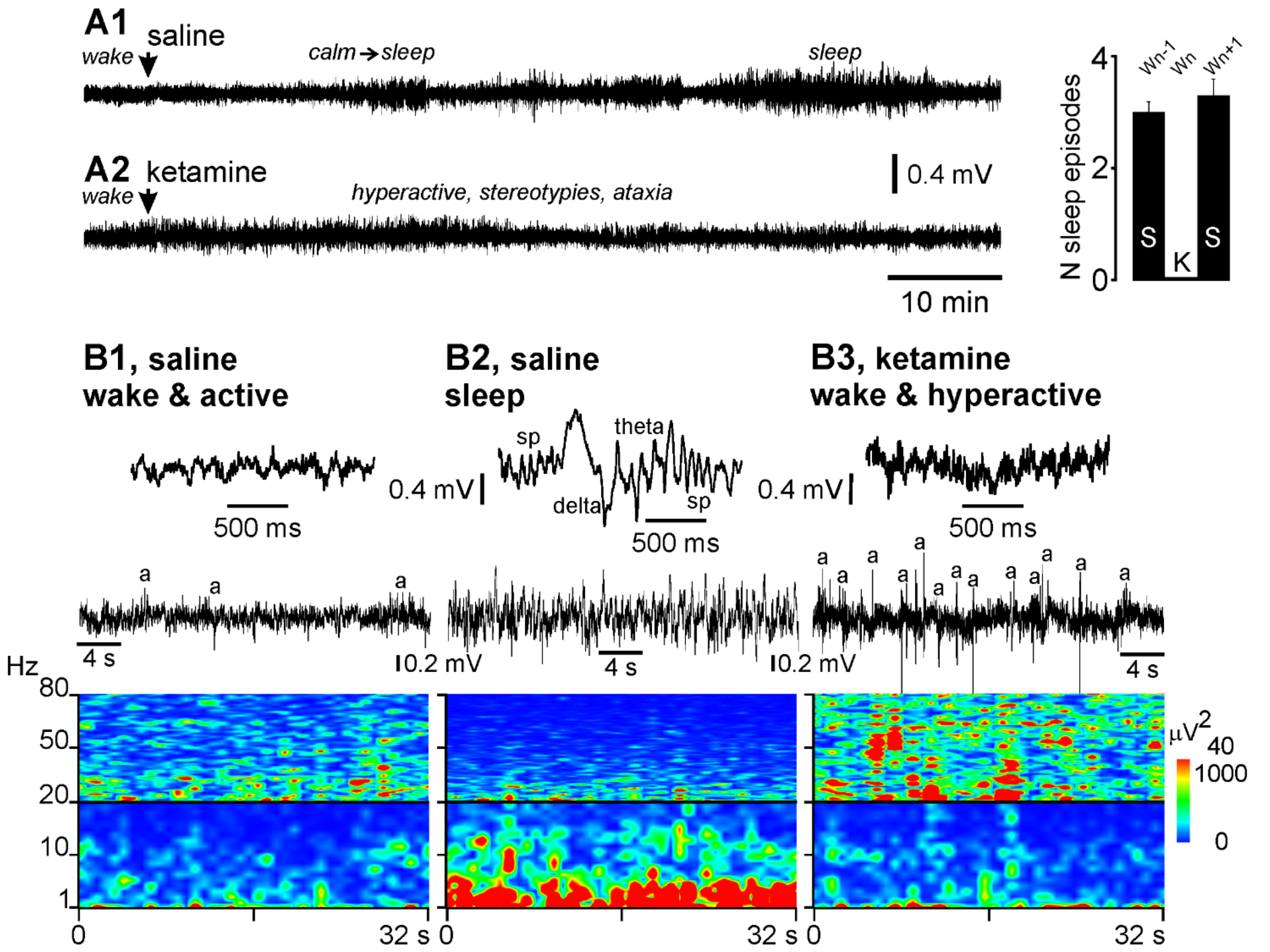


**Figure S1: Ketamine prevents the occurrence of natural sleep episodes in free-behaving rats**.
**(A1-A2)** Two typical 80-min recording sessions (cortical EEG, bandpass: 1-800 Hz, recording cable-induced artifacts withdrawn) performed in a freely moving rat with one week interval under the saline (A1) and ketamine (A2) conditions. Saline (1 ml/kg) or ketamine (2.5 mg/kg, 1 ml/kg) was subcutaneously injected a few min after the beginning of the recording session (arrow). At the very beginning of the session, the rat is usually active with a cortical EEG displaying, prominently, low-amplitude (<0.2 mV) and fast (frequency band > 17 Hz) oscillations (B1). Under control (saline) condition, the rat usually becomes calm (quiet immobile wakefulness) then, after toileting, goes into slow-wave sleep episodes. Non-REM sleep episodes were identified on the basis of the occurrence of delta (1-4 Hz)-, theta (5-9 Hz)- and spindle (sp, 10-16 Hz)-frequency oscillations (B2). Under the ketamine condition, the rat becomes quickly hyperactive (erratic behavior with stereotypies and ataxia) associated with a cortical EEG characterized by an abnormally excessive and persistent amplification of ongoing beta- and gamma-frequency (18-29 and 30-80 Hz, respectively) oscillations (B3) with a partial recovery at the end of the recording session. No one sleep episode was identified during the 80-min recording session. The histogram on the right illustrates the number of non-REM sleep episodes during the 80-min recording session under the control (S or saline; N = 3.0±0.2, 18 recordings sessions), one week before (Wn-1) the ketamine challenge, or under the ketamine (K, 0.0±0.0, 15 recordings sessions) conditions. One week later (Wn+1), the ketamine-treated rats retrieve their sleep episodes under the saline condition (n = 3.3±0.3, 11 recording sessions). **(B1-B3)** Top traces: short bouts of desynchronized (during wake state, B1) and synchronized (during non-REM sleep, B2) cortical EEG recorded under saline condition, and of “hyper-desynchronized” cortical EEG under ketamine condition (B3, 7 minutes after injection). Middle traces: 32-s bouts of wake-related desynchronized (during wake state, B1) and non-REM sleep-related synchronized (B2) cortical EEG recorded under saline condition, and of ketamine-induced “hyper-desynchronized” cortical EEG (B3, 7 minutes after injection). Under the ketamine condition (B3), the rat behavioral hyperactivity generated many artifacts (a) in the cortical EEG due to intense electromyographic activities and cable artifacts. Bottom: Time-frequency spectral analysis (resolution: 0.03 Hz, hamming, 50% overlap) of a 32-s recording episode for each condition.

**Figure S2: Sleep-like activity patterns under the analgesic pentobarbital sedation.**

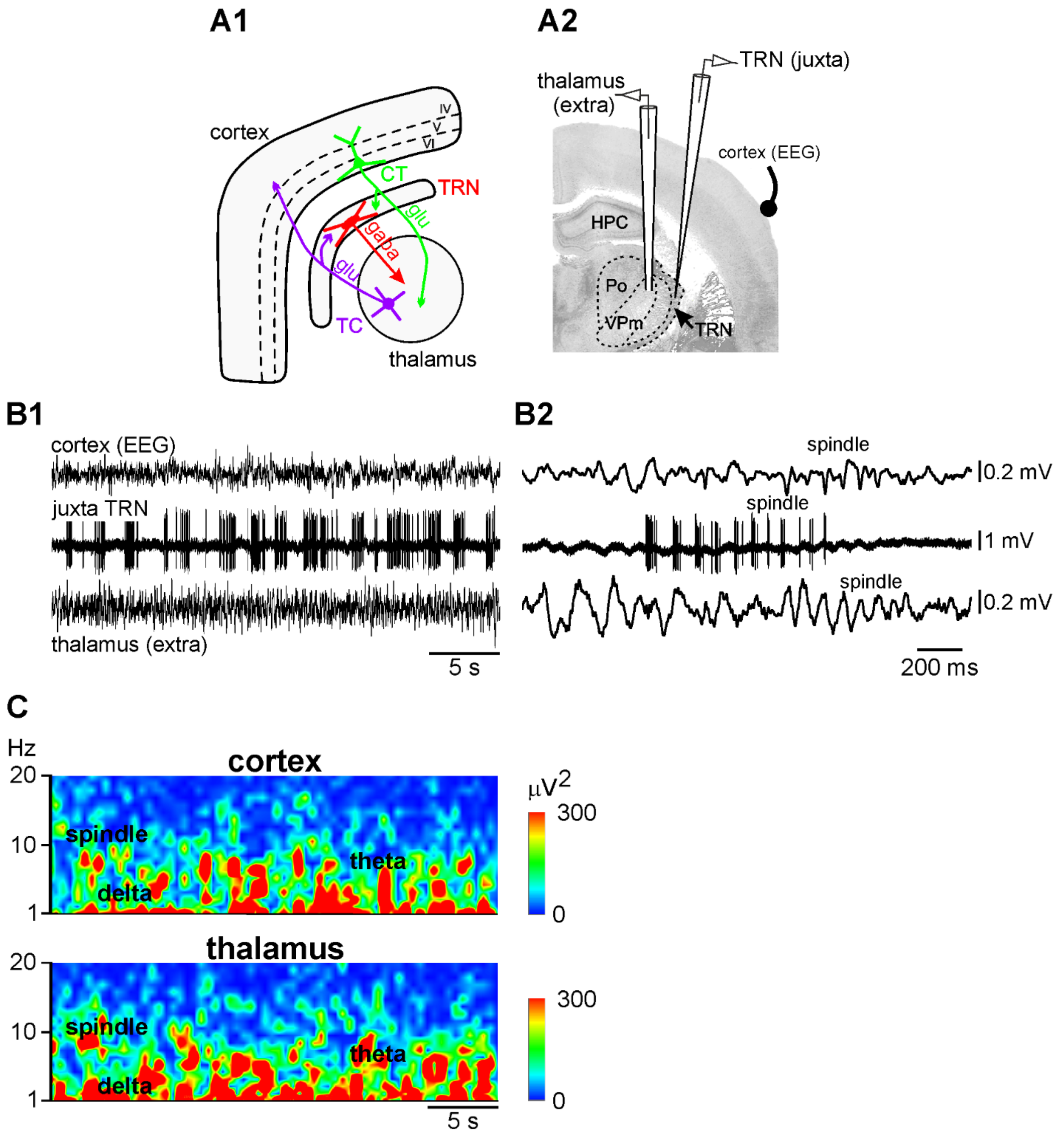


**(A1)** The hodology of the layer VI CT-TRN-TC circuit, the common circuit of both first- and higher-order TC systems and the leading circuit in the generation of spindles. **(A2)** Location of the cortical (EEG), thalamic (extracellular configuration), and of the TRN (juxtacellular configuration) electrodes in the somatosensory system. **(B1)** A 32-s trace recorded in a pentobarbital sedated rat. Both the cortical EEG and the thalamic extracellular activities are prominently synchronized. During this synchronized state, the TRN cell fires rhythmic high-frequency bursts of APs at the delta- (1-4 Hz), theta- (5-9 Hz) and spindle- (10-16 Hz) frequency bands. **(B2)** A short-lasting trace showing spindle network (Cx EEG and thalamus extracellular) and cellular (TRN) activities. **(C)** Time-frequency spectral analysis of the 32-s cortical and thalamic records (resolution: 1 Hz, hamming, 50% overlap) of the B1 trace.

**Figure S3: Ketamine significantly decreases the power of spindles and delta-frequency oscillations and increases that of gamma- and higher-frequency oscillations in the somatosensory thalamocortical system.**

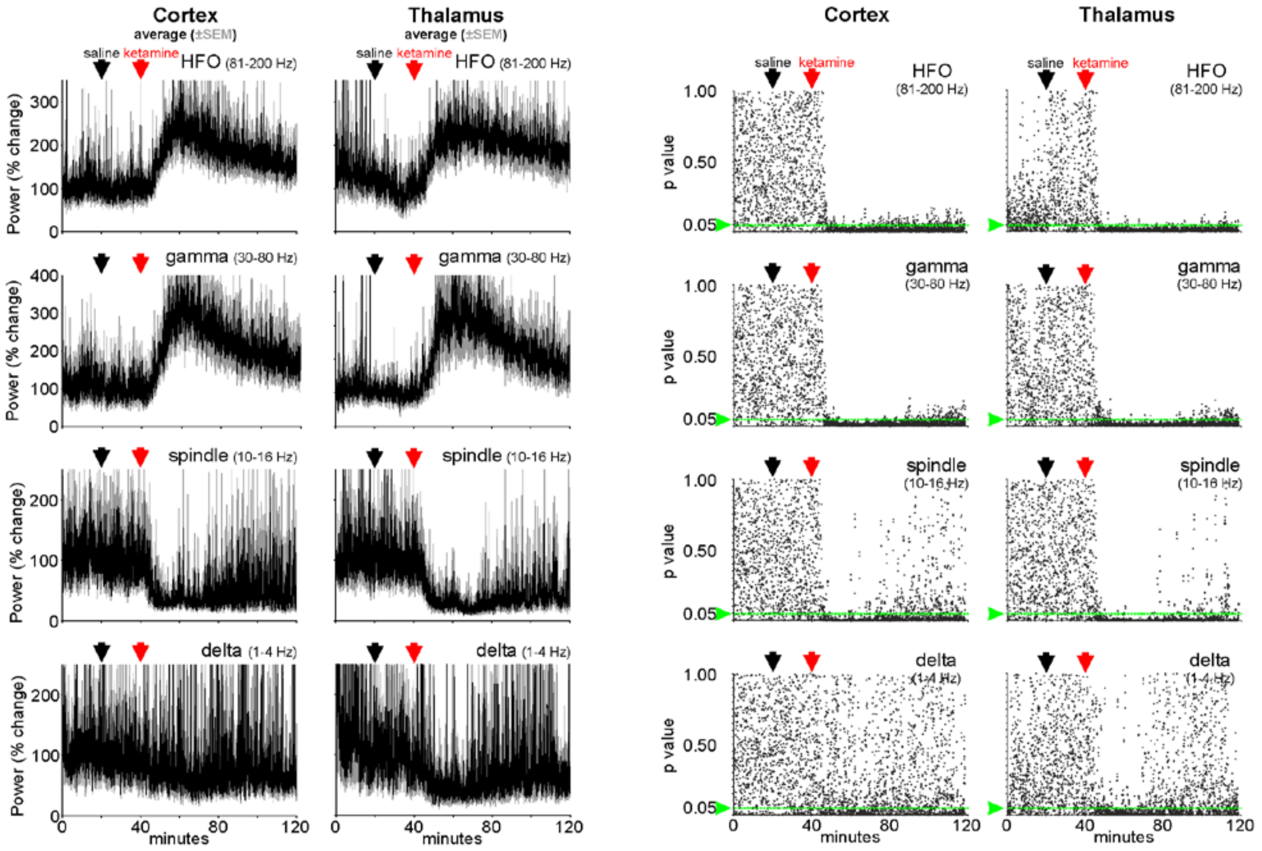


**(left panel)** Time course of the power (% change relative to the saline condition, grand average from 6 rats with simultaneous cortical and thalamic recordings) of neural oscillations before and after the subcutaneous administrations of saline (at 20 min) and ketamine (at 40 min). **(right panel)** Each dot is a Student's paired t-Test (one test ketamine relative to saline (100%) every 2 seconds). The 100% corresponds to the average of all the values (n=300 per rat) obtained during the 10 min period that precedes the ketamine administration. The p-value 0.05 is indicated by the green line. The smaller the p-value, the higher the significance. HFO, high-frequency oscillations.

### S4: MK-801 mimics the ketamine effects better than physostigmine and apomorphine

There is accumulating evidence that ketamine acts at diverse receptors other than NMDA receptors, including dopaminergic and cholinergic receptors (1, 2). Therefore, one may wonder whether the acute ketamine effects, observed under our experimental conditions, were specifically due to a blockade of NMDA receptors. In an attempt to address this issue, the sleep-related cortical EEG oscillations were recorded following the subcutaneous administration of either MK-801 (dizocilpine, 0.1 mg/kg), a more specific non-competitive antagonist of NMDA receptors, the cholinesterase inhibitor physostigmine (0.5 mg/kg), or the competitive dopamine D2 receptor agonist apomorphine (1 mg/kg). MK-801 was expected to reproduce the ketamine effect (3), physostigmine to promote REM sleep (4) and a desynchronized cortical EEG (5, 6), and apomorphine to affect the EEG oscillations (7, 8).

Like ketamine, MK-801 consistently decreased the power of ongoing spindles and substantially increased that of gamma- and higher-frequency TC network oscillations (FigS4a, FigS4b). No recovery of the MK-801 effects was observed 80 minutes after the administration. In contrast to ketamine, MK-801 did not significantly change the power of delta-frequency oscillations (FigS4b), very likely the result of a dose effect (9).

Unlike ketamine, physostigmine or apomorphine significantly decreased the power of all frequency bands in TC network oscillations (FigS4a, FigS4b). The pharmacokinetics of these two drugs present a plateau-like curve, beginning at about 10 and 20 minutes and starting to recover at the end of the recording session. In juxtacellularly recorded TRN neurons, physostigmine (n = 2 TRN cells from 2 rats) decreased the density of hfBursts and increased that of sAPs, and apomorphine (n = 2 TRN cells from 2 rats) decreased, in a similar way, the density of both sAPs and hfBursts (FigS4c).

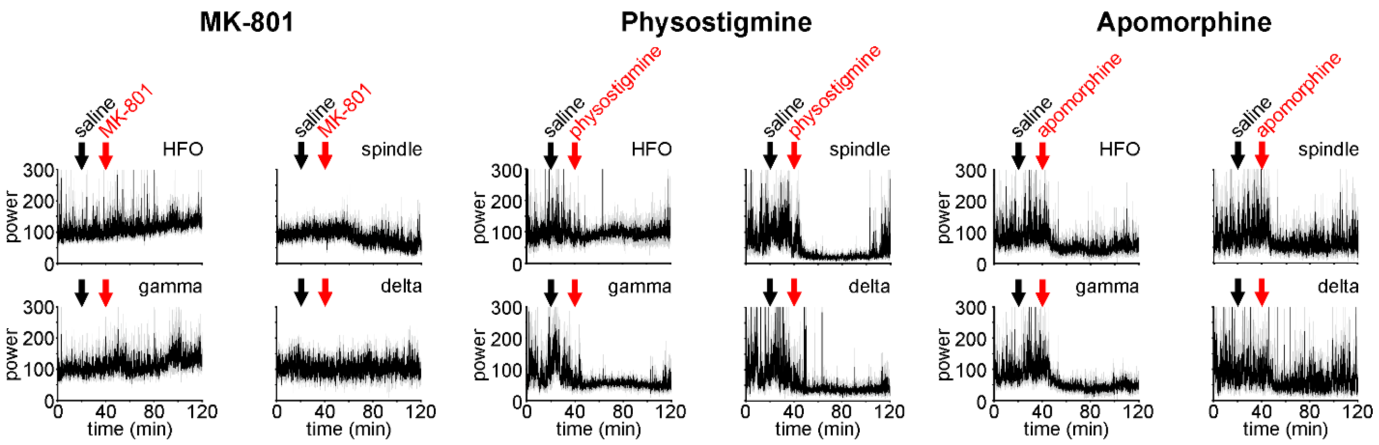


**Figure S4a: MK-801 mimics the ketamine effects better than physostigmine and apomorphine.** Time course of the power (% change relative to the saline condition, mean (in black) ±SEM (in grey), 4 rats per condition, each rat being its own control) of cortical EEG oscillations (all frequency bands) before and after the subcutaneous administrations of saline (at 20 min) and (at 40 min) either MK-801 (0.1 mg/kg), physostigmine (0.5 mg/kg), or apomorphine (1 mg/kg).

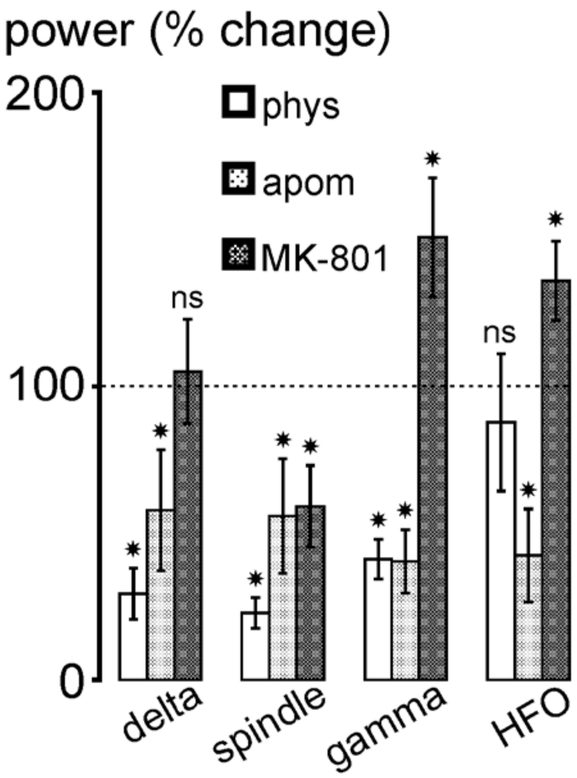


**Figure S4b: MK-801 mimics the ketamine effects better than physostigmine and apomorphine** The histogram illustrates the drug-induced percent changes (at 20-60 min postinjection, relative to the saline condition, each animal being its own control) in power of the four frequency bands in the cortical EEG (4 rats per drug; mean±SEM). Student t-test: (*) p< 0.001; ns, not significant.

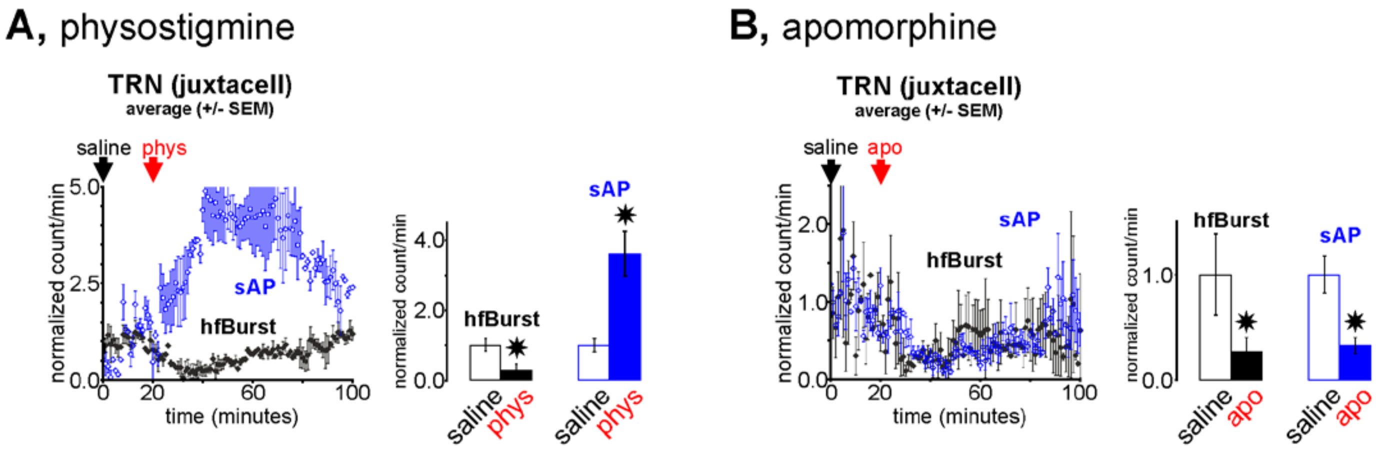


**Figure S4c: Physostigmine or apomorphine decreases the hfBurst density in juxtacellularly recorded thalamic reticular nucleus neurons.**
The density (number per minute, ±SEM, 2 TRN cells from 2 rats) of hfBursts and of sAPs under the saline (A or B), physostigmine (A) or apomorphine (B) conditions. In the histograms, the normalized values are from the time period of 20-60 min postinjection (phys or apo), that is, 40-80 min in the charts. Note that, like ketamine, physostigmine increases the sAP density and decreases the hfBurst density.

**Figure S5: Ketamine switches the firing pattern of thalamic reticular nucleus (TRN) neurons from the hfBurst mode to the sAP mode.**

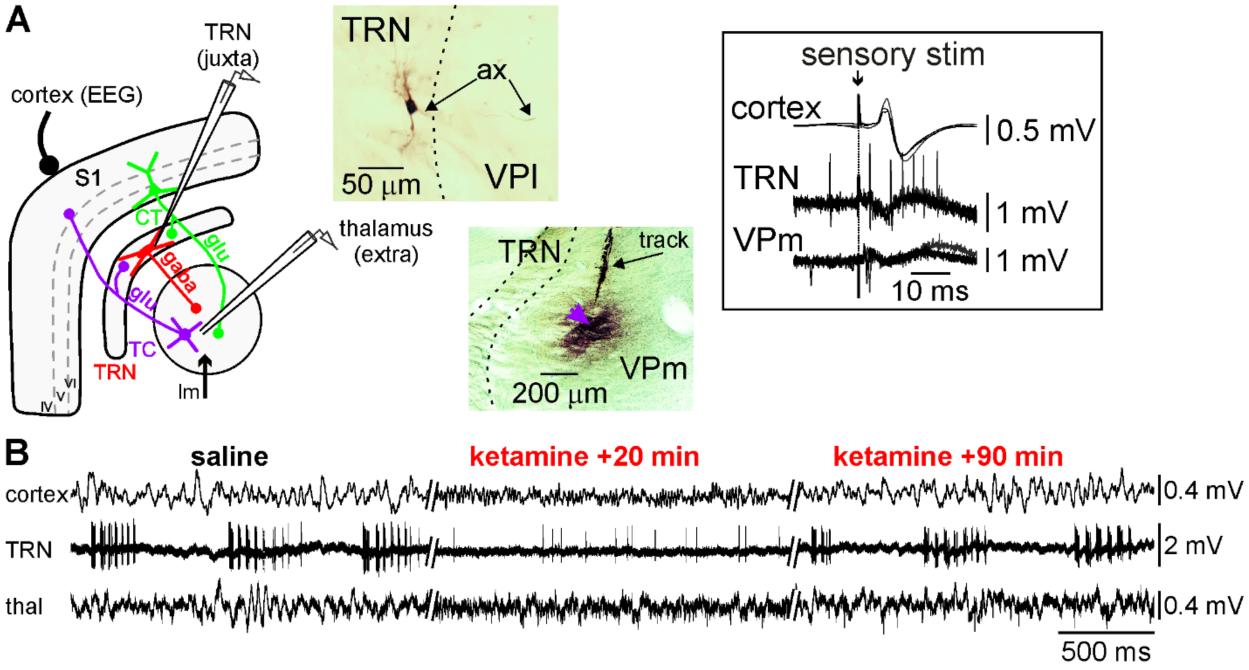


**(A)** Experimental design showing the location of the recording glass micropipettes, a sharp one (tip diameter ~1µm) to record juxtacellularly (juxta) a single thalamic reticular nucleus (TRN) neuron, a semi-micropipette (tip diameter: 5-7 µm) to record the extracellular activities of a subpopulation of TC neurons in the somatosensory system (VPm, medial part of the ventral posterior nucleus). These intrathalamic recordings are done along with the cortical EEG of the related primary somatosensory cortex (S1). Is also shown, the hodology of the somatosensory 3-neuron layer VI CT-TRN-TC circuit, the principal leading cicuit in the generation of spindles. In the somatosensory system, the lemniscal (lm) input being the principal prethalamic input of the VPm. The corticothalamic (CT) and TC neurons are glutamatergic while the TRN neuron is GABAergic. At the end of the recording session, the location and the structure of the recorded neurons are labelled with the neuronal tracer Neurobiotine. The top microphotograph shows part of the somatodendritic complex and the main axon (ax) of a juxtacellularly recorded TRN neuron; the bottom microphotograph shows the extracellular labelling (methyl green counterstaining) of the recording site and electrode track in the VPm, the head arrow indicating the location of the extracellular recording site. In the frame is shown the functional identification of the recorded somatosensory neurons at the beginning of the recording session, that is, short-latency sensory-evoked activities simultaneously recorded in the cortical EEG and in thalamic relay (VPm) and reticular (TRN) neurons. **(B)** Typical simultaneous recordings of the S1 cortex (EEG), the somatosensory TRN (single-unit juxtacellular configuration) and related thalamus (extracellular configuration). Under the saline (control) condition, both the cortex and the thalamus exhibit a synchronized state, characterized by the occurrence of low-frequency (1-16 Hz) oscillations, including spindles, and the TRN cell exhibits three series of rhythmic robust high-frequency bursts of action potentials (300-500 APs/s). Under the ketamine condition (here, 15 minutes post-ketamine injection), the TC system displays a more desynchronized state, characterized by the prominent occurrence of faster activities (>16 Hz), which include gamma-frequency oscillations, and the TRN cell fires more in the single AP mode than in the burst mode. Ninety minutes after the subcutaneous administration of low-dose ketamine, the sleep state of the TC system is back in the CT-TRN-TC system.

**Figure S6: Ketamine decreases the high-frequency burst (hfBurst) density and increases the single AP (sAP) density in extracellularly recorded thalamocortical (TC) neurons.**

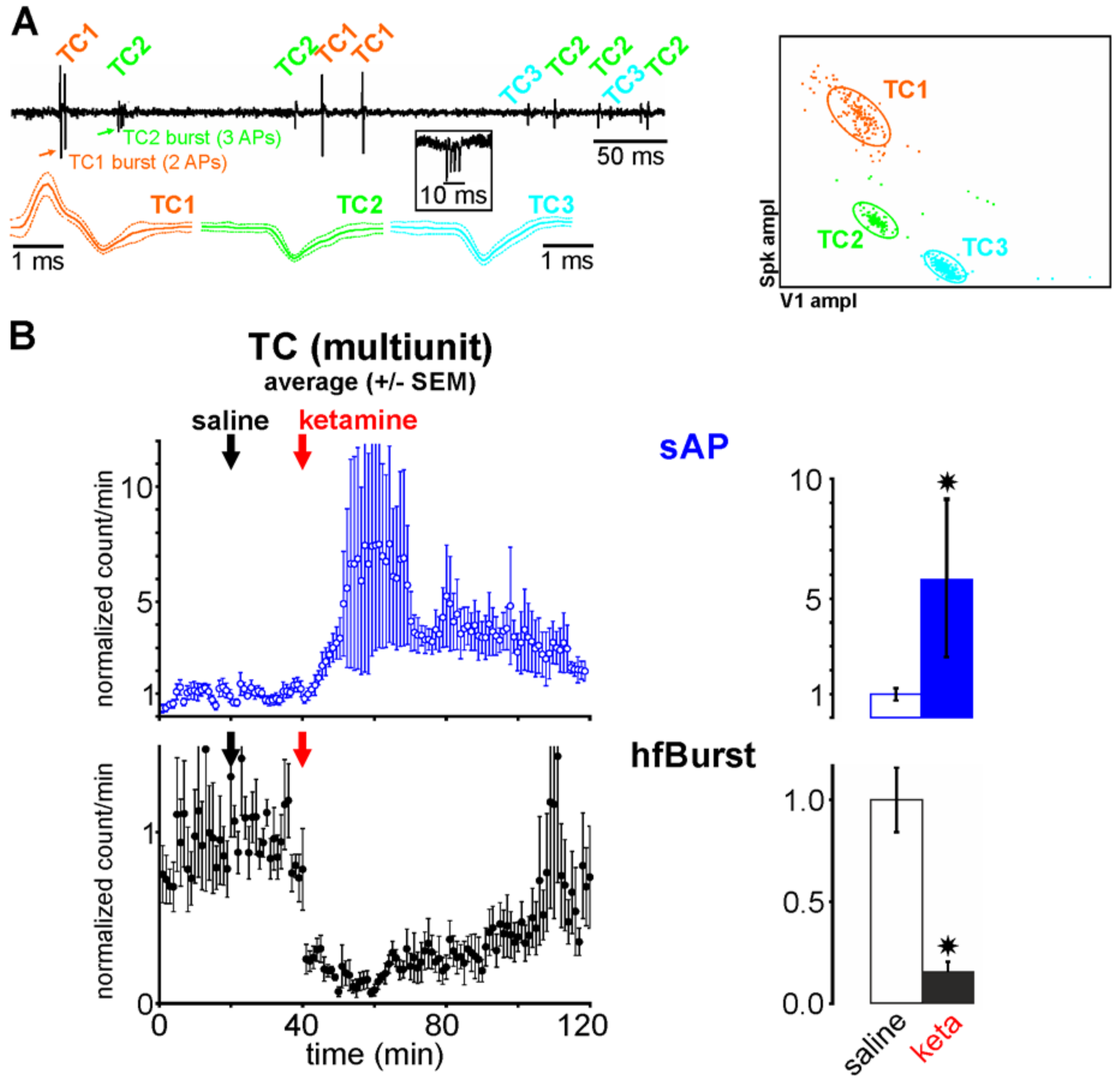


**(A)** A typical example of a spike sorting of 3 TC cells (TC1, TC2 and TC3) from an extracellular multiunit recording. In the recording bout (high-pass filter cut at 100 Hz), the three TC cells are visually well distinguishable, TC1 exhibiting a hfBurst of 2 APs then sAPs, TC2 a hfBurst of 3 APs then sAPs, and TC3 only sAPs. A typical extracellular hfBurst is shown in the frame. The mean±SD of 50 APs of the three detected TC neurons are shown. Three clusters are well distinguishable on the basis of the amplitude of the spike and valley components (spk ampl and V1 ampl, respectively) of the APs. **(B)** Grand average (±SEM, N=16, from 6 rats) of the relative changes of the density (normalized count per minute, 1 being the control value under the saline condition) of the sAPs and of the hfBursts. On the right, the histograms show the average values corresponding to 20-40 minutes ketamine postinjection. Asterisk when significant (paired t-test, $p<0.01$).

**Figure S7: Ketamine increases the firing frequency band similarly in two juxtacellularly recorded nearby TC neurons.**

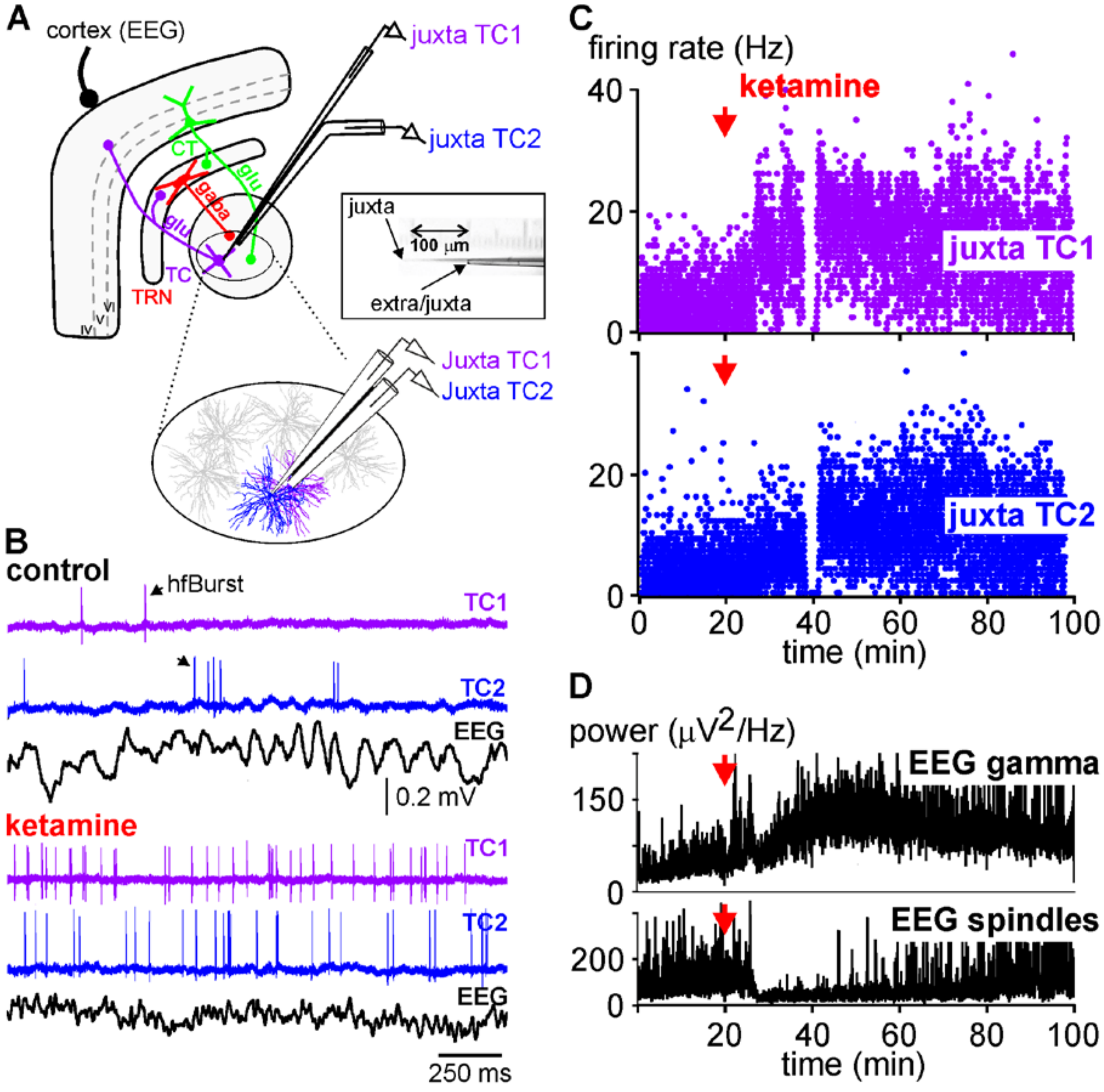


**(A)** Experimental design showing the location of the recording EEG electrode on the primary somatosensory cortex and of the combined juxtacellular (tip diameter: 1 µm) and extracellular (tip diameter: 5 µm) glass micropipettes (inter-tip distance: 100 µm, see microphotograph in the frame). In this experiment, the latter electrodes are located in the posterior group of the thalamus (the upper lip being the receptive field for the juxtacellular micropipette). **(B)** Each of the two combined micropipettes records juxtacellularly a single TC neuron. Under the control (saline) condition, the cortical EEG displays sleep oscillations, including spindles, whereas both TC1 and TC2 neurons fire sAPs and hfBursts. Four to 5 minutes after a subcutaneous administration of low-dose ketamine (2.5 mg/kg), the cortical EEG exhibits less spindles/slower oscillations and more higher-frequency (>16 Hz) oscillations, including especially gamma oscillations, and the two adjacent TC neurons fire more sAPs than hfBursts. **(C)** Ratemeter of the simultaneously juxtacellularly recorded TC1 and TC2 neurons under saline then ketamine conditions. Each dot corresponds to the number of inter-AP intervals per second. At 40 minutes, no available data for a short while because of the sudden occurrence of artifacts that prevented accurate AP detection. **(D)** Time course of the power of gamma oscillations (top) and of spindles (bottom) recorded simultaneously in the somatosensory cortex before and after subcutaneous administrations of saline and ketamine (at 0 and 20 min, respectively).

**Figure S8: Time relationship between the APs of a single TC neurons and the related juxtacellular, extracellular and cortical gamma waves.**

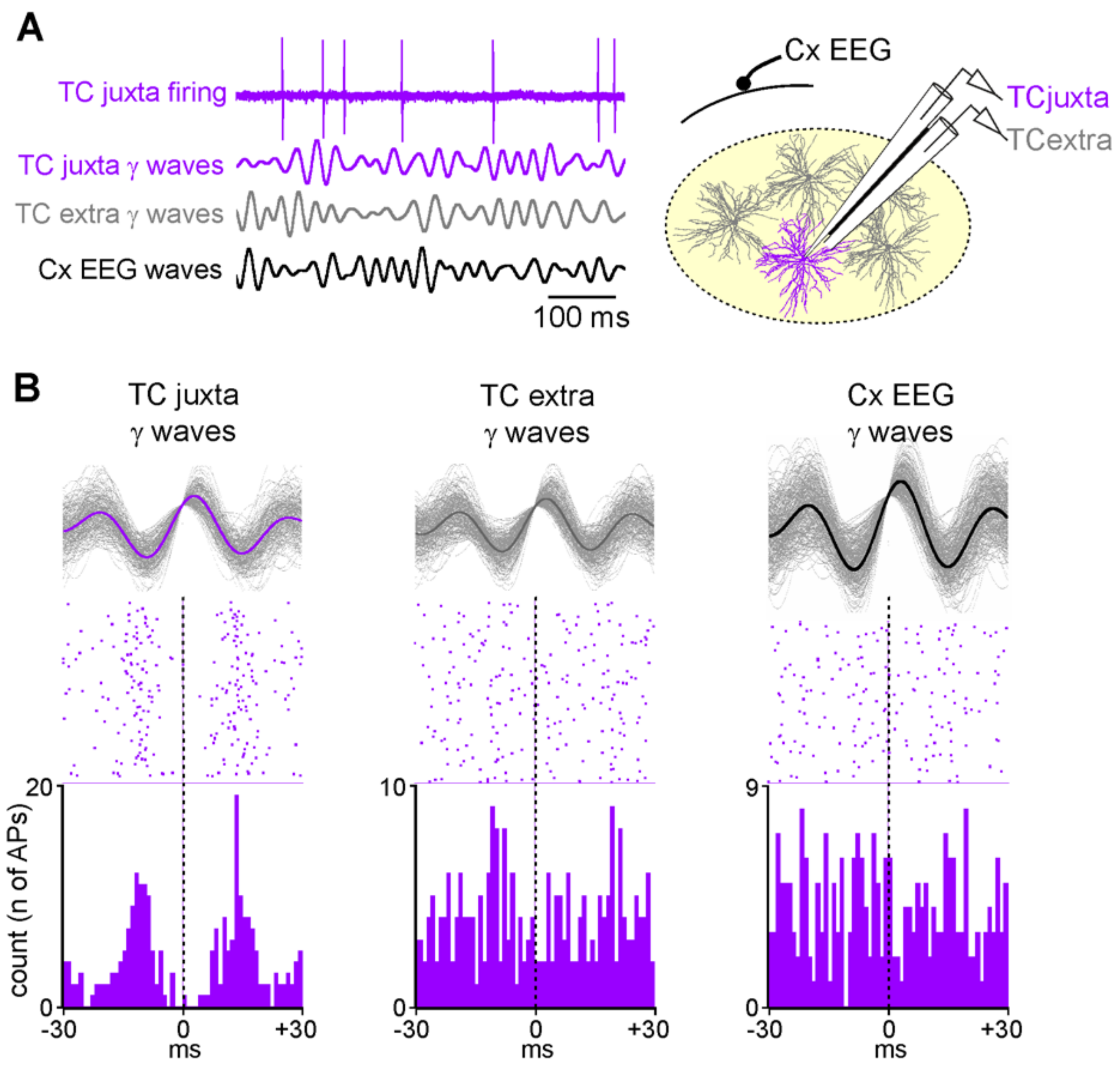


**(A)** Simultaneous dual juxtacellular-extracellular recording of a single TC neuron along with the EEG of the related cortex of the somatosensory system. A schematic drawing of the experimental design is shown on the right. From top to bottom: Juxtacellular firing of a single TC neuron (bandpass: 1-6000 Hz); juxtacellular TC gamma oscillations (25-55 Hz); extracellular (100 μm distant from the neuron) TC gamma oscillations (25-55 Hz); cortical EEG gamma oscillations (25-55 Hz). **(B)** Peri-event (gamma wave) time histogram (1-ms resolution) of the TC firing (cumulative count) under the ketamine condition. Every gamma wave (TC juxta, TC extra (inter-tip distance = 100 μm, see drawing), and Cx EEG) is an average of 100 filtered (25-55 Hz) individual gamma (γ) waves. Time “0” corresponds to the time at which gamma waves were detected. Two hundreds sweeps, each dot representing a detected AP.

### S9: Ketamine strengthens the gamma-frequency band TRN-TC connectivity

The present findings support the hypothesis that TC and TRN neurons are endowed with intrinsic and synaptic properties to generate TRN-TC gamma-frequency oscillations (1). If it is true that the TRN is the pacemaker of thalamic gamma-frequency oscillations, then it is expected that the TRN-TC gamma connectivity increases after the ketamine administration.

**Methods and Materials:** To investigate changes in the strength of gamma interactions after ketamine application within the CT-TRN-TC network, all signals (ECoG, TRN juxta and TC extra) were filtered in the gamma range (25-55Hz). Direct interaction strength between 2 different sites *i* and *j* was given by partial correlation coefficient $\Pi_{ij}$ (2). Unlike usual correlation coefficient, partial correlation coefficient considers only the "direct interaction" between the sites, i.e. this correlation is free from the influence of any other site in the network. Partial correlation coefficients were computed via the following formula (3):

$$\Pi_{ij} = -\frac{\Upsilon_{ij}}{\sqrt{\Upsilon_{ii}\Upsilon_{jj}}}$$

Where $\Upsilon$ stands for the inverse covariance matrix $\Sigma^{-1}$ between two signals. The coefficients of the covariance matrix were taken as the amplitudes of the central peaks from the cross-correlograms of typical 400ms-epochs from signals recorded at the corresponding sites. The resulting partial correlation coefficients for CT-TRN, CT-TC, and TRN-TC interactions were compared under both control and ketamine conditions using paired t-test.

**Results**:

We assessed this gamma connectivity in the somatosensory layer VI CT-TRN-TC circuit. Under control condition, the strength of gamma interactions, given by partial correlation coefficient (see methods), was higher between TRN and TC sites than for CT-TRN and CT-TC interactions (FigS9). This fact was also reflected by a high central peak in the average cross-correlogram of TRN and TC signals. In contrast to this, average CT-TRN and CT-TC cross-correlograms did not exhibit a well-defined central peak. When ketamine was applied, the strength of TRN-TC gamma interactions was significantly increased compared to control condition (paired t-test, $p<0.001$, $n=11$), resulting in a higher partial correlation coefficient and a higher peak in the average cross-correlogram. CT-TRN and CT-TC cross-correlations also increased and a well-distinguishable central peak appeared in their cross-correlograms (Fig6). However, the strength of CT-TRN and CT-TC interactions given by partial correlation coefficients did not change significantly compared to control condition (paired t-test, $p>0.4$), suggesting that increase in these cross-correlations is artificial and cannot be attributed to the changes in the direct strength of their gamma connectivity but should be rather explained by other mechanisms (influence of other network sites, volume conduction, etc).

1. Pinault D, Deschênes M (1992): Voltage-dependent 40-Hz oscillations in rat reticular thalamic neurons in vivo. *Neuroscience*. 51:245-258.
2. Marrelec G, Krainik A, Duffau H, Pelegrini-Issac M, Lehericy S, Doyon J, et al. (2006): Partial correlation for functional brain interactivity investigation in functional MRI. *Neuroimage*. 32:228-237.
3. Whittaker J (2009): *Graphical Models in Applied Multivariate Statistics*. 1st ed. Chichester England; New York: Wiley.

### Figure S10: Clozapine prevents the ketamine effects on sleep TC oscillations

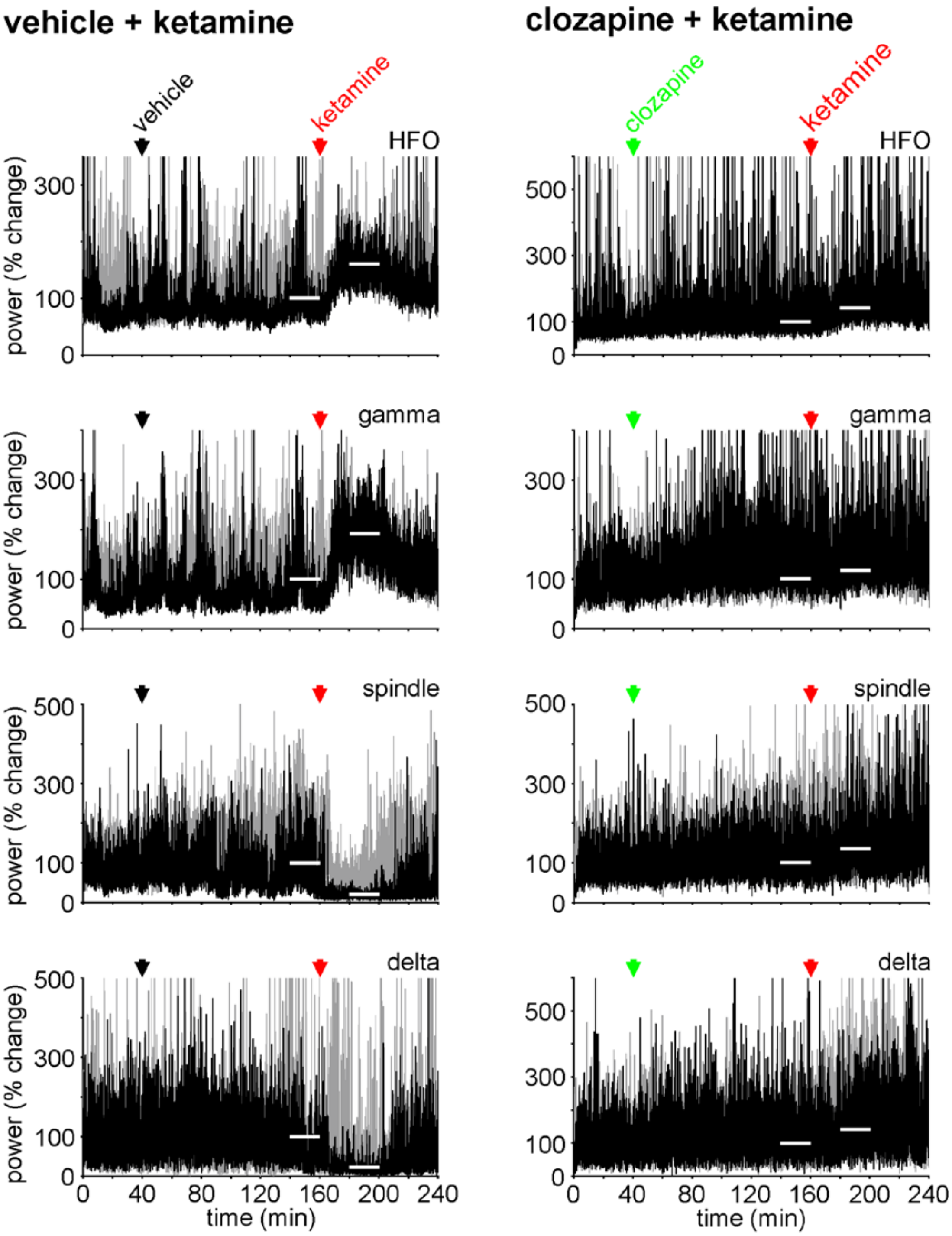


Each chart shows, for a given frequency band oscillation, the time course (1 FFT value/2 seconds) of the drug effects (% change in power) on the cortical EEG in 2 individuals (grey and black) during a 240 min recording session, the ketamine challenge being done at 160 minutes under the control (left panel) and clozapine (right panel) conditions. The horizontal white bars indicate the time periods used for statistical comparisons (paired t-test). **Left panel**: Ketamine (2.5 mg/kg) was administered 120 minutes after the vehicle (NaCl/HCl 0.1N) administration (subcutaneous, 1 ml/kg). **Right panel**: Ketamine (2.5 mg/kg) was administered 120 minutes after the clozapine (dissolved in NaCl/HCl 0.1N) administration (subcutaneous, 5 mg/kg, 1 ml/kg). HFO, high-frequency oscillations (81-200 Hz).

### S11: Downsides and upsides

From a pharmacological viewpoint, it is hazardous to draw firm conclusions as all experiments designed to assess the effects of any tested drug were conducted in the pentobarbital-sedated rat. Although that the pentobarbital may somehow have interacted with the action of any of the drugs tested, the sleep model offers the advantage to assess the acute ketamine effects simultaneously on the whole spectrum of neural oscillations, which are translational electro-biomarkers and potential endophenotypes of a psychosis transition state.

From a conceptual and translational point of view, it is amazing to see that the neurophysiological effects of the NMDA receptor antagonists, observed under the present experimental conditions, are fairly similar to those recorded in awake, naturally behaving rats (1-3). Also, the ketamine-induced changes in rodent brain EEG oscillations are reminiscent of those observed in at-risk mental state individuals (4, 5) and during the first-episode of schizophrenia (6, 7). So what does the acute ketamine model indeed model? First of all, it is worth saying that this model does not consider the genetic-neurodevelopmental and anatomical dimension, and the decompensatory mechanisms of psychotic disorders. The acute ketamine rodent or human model is thought to model acute forms of psychosis and better early than chronic schizophrenia (8). Ketamine alone initiates NMDA receptor-related primary and cascade mechanisms, which seem sufficient to cause a transient psychosis-relevant state. In the present preclinical investigation, ketamine elicited a transition from a physiologically “normal” state to a psychosis-relevant state. On the other hand, in humans, the natural transition to psychosis emerges from an already disturbed state (or at-risk mental state), which implicate multifactorial, interactive etio-pathophysiological mechanisms involving environment, socio-culture, genes, pro-inflammation, risk factors, dopaminergic, GABAergic and glutamatergic neurotransmissions, and functional dysconnectivity between cortical and subcortical highly-distributed systems, including thalamus-related circuits. In spite of the disputable validity of the acute ketamine model, the present study may help to understand, at least in the highly-distributed TC systems, the cell-to-network disturbances occurring during the conversion from a physiological to a pathological, psychosis-relevant state (Fig8). Cortical EEG oscillations are reliable bio-markers of thalamic activities as both the cortex and the thalamus work together. In the present study, it is shown that it is possible, based on the available knowledge, to predict the likely thalamic cell-to-network correlates. Therefore, the present findings provide a translational added value to the present conceptual approach with some construct validity. However, it should also be mentioned that, in patients with psychosis, the natural EEG disturbances (reduction in spindle activity, increase in gamma power, etc.) are subtler than the ketamine-induced changes in brain activities.
